# Gapless *indica* rice genome reveals synergistic effects of active transposable elements and segmental duplications that promote rice genome evolution

**DOI:** 10.1101/2020.12.24.424264

**Authors:** Kui Li, Wenkai Jiang, Yuanyuan Hui, Mengjuan Kong, Li-Zhi Gao, Pengfu Li, Shan Lu

## Abstract

The ultimate goal of genome assembly is a high-accuracy gapless genome. Here we report a new assembly pipeline which we have used to produce a gapless genome for the *indica* rice cultivar Minghui 63. The 395.82 Mb final assembly is composed of 12 contigs with a contig N50 size of 31.82 Mb. All chromosomes are now gapless, with each chromosome represented by a single contig. This is the first gapless genome assembly achieved for higher plants or animals. BUSCO evaluation showed that gene regions of our assembly have higher completeness than the current rice reference genome (IRGSP-1.0). Compared with *japonica* rice, *indica* has more transposable elements (TEs) and segmental duplications (SDs), the latter of which produce many duplicated genes that can affect plant traits through dose effect or sub-/neo-functionalization. The insertion of TEs can also affect the expression of duplicated genes, which may drive evolution of these genes. We also found the expansion of *NBS-LRR* disease resistance genes and *cZOGT* growth-related genes in SDs, suggesting that SDs contribute to the adaptative evolution of rice disease resistance and developmental processes. Our findings suggest that active TEs and SDs together provide synergistic effects to promote rice genome evolution.

## Introduction

Repeat sequences, such as transposable elements (TEs), centromeres, and segmental duplications (SDs), represent the most challenging genomic regions to assemble (Alkan et al., 2011; Bailey et al., 2001; Elert, 2014; Liu et al., 2020a; Michael and VanBuren, 2015; Numanagic et al., 2018). However, these regions can play major roles in the genome evolution of a lineage. For example, TEs participate in shaping gene expression patterns and the formation of genome structure. This is especially true for long terminal repeat retrotransposons (LTR-RTs), which are the most prevalent repeats in plant genomes, and their proliferation could result in genome bloating (Elert, 2014). Other repeat regions, such as centromeres, are of central importance to chromosome stability (Mackinnon et al., 2013). SDs are similarly influential on genome structure due to their production of numerous repeat genes, and thus serve as hotbeds for genomic rearrangements followed by gene innovation and rapid gain-of-function adaptations (Han et al., 2009; Marques-Bonet et al., 2009; Zhang et al., 1998).

With the development of long-read sequencing technology, the continuity of genome assembly has been greatly increased by orders of magnitude. Some recently developed technologies such as binano (Jiao et al., 2017), HIC (Dudchenko et al., 2017), and 10x Genomics (Mostovoy et al., 2016; Yeo et al., 2018) have further improved the continuity of assembly. The genomes of rice (Du et al., 2017), maize (Liu et al., 2020a), and rose (Raymond et al., 2018), for example, have only a few unresolved gaps. However, the ultimate goal of assembly, *i.e.*, a highly accurate gapless genome, has yet to be achieved. High-fidelity (HiFi) sequence reads, generated via circular consensus sequencing (CCS), are the first data type that has advantages in both read lengths (greater than 10 kb) and accuracy (greater than 99%). In this method, subreads from a single polymerase read are computationally combined via the CCS algorithm to create a HiFi consensus read (Logsdon et al., 2020). This produces high-quality reads without overlapping and error correction, which may introduce errors caused by incorrect overlaps (especially for repeat sequence regions), making it possible to obtain high-quality genome assemblies.

Since rice (*Oryza sativa* L.) is a staple crop consumed by more than half of the global population (Elert, 2014), its genome is of broad interest for studies from the relationship and impacts of evolution on plant traits to the selection for high yield or disease resistance to maximize production. *Indica* rice is the most widely produced rice subspecies due to several desirable traits, and interestingly, has a substantially larger genome than another cultivated subspecies *japonica* (Du et al., 2017). Recent studies have shown that TEs, centromeres, and SDs all have strong evolutionary effects on the genome (Elert, 2014; Han et al., 2009; Mackinnon et al., 2013; Marques-Bonet et al., 2009; Zhang et al., 1998). However, the prevalence of these repeat sequences in plant genomes also hinders the assembly of high-quality genomes, for which current sequencing technologies and algorithms have considerable difficulties in chromosome-level assemblies without gaps. Here, with the help of other genome research methods, such as genetic maps (Catchen et al., 2020), HIC (Dudchenko et al., 2017), or alignment with genome sequences of related species, we developed a feasible approach to achieve the gapless assembly of the *indica* rice cultivar Minghui 63 based on HiFi reads. Further, we used our gapless genome to prove the hypothesis that recent TEs and duplicated genes in *indica* rice provide synergistic effects on genome evolution that impact adaptative traits. Our results indicate that *indica* rice has many SDs that result in gene duplications necessary for disease resistance and biomass accumulation. We also found a preponderance of active TE insertions in the proximity of the duplicated genes with low expression levels, while a few highly expressed duplicated genes appeared to have undergone sub- or neo-functionalization.

## Results

### Gapless assembly of the rice genome and feature annotation

We used PacBio HiFi reads from *indica* rice variety Minghui 63 to reassemble the genome. PacBio HiFi reads provide long fragment reads with 99% accuracy, which avoid the necessity of error correction before assembly. This is particularly useful for the assembly of high-repeat areas such as centromeres, telomeres, TEs, and SDs, in which read alignments are often collapsed during error corrections. Briefly, 25.3 Gb HiFi reads from the National Center for Biotechnology Information (NCBI) Sequence Read Archive database (SRX6957825) were used for preliminary assembly using pb-assembly (Wenger et al., 2019) (Fig 1A, Table S1). Then, the reference genome of *indica* rice (MH63RS2) (Zhang et al., 2016) was used to anchor contigs onto chromosomes (Fig. 1B). With the mapping for each contig location (Table S2), every chromosome was represented by a list of contig paths (encoding the graph for each contig and represented by a list of unitigs) and gaps between them. If two contigs were adjacent, they were connected without a gap. After this step, there were only 25 gaps remained between contig paths. We further reconstructed the string graph using edges (sg_edges_list) and overlap information (pread.m4) generated by pb-assembly, found all the paths between each gap, and then selected the path with the most overlaps for each gap. Most of the paths were simple paths, except for a few complex paths spanning multiple repeat regions (Fig. 1C). Through the above steps, 24 gap areas were filled and one gap was removed for two flanking contigs becoming adjacent after removing possible assembly errors. We obtained a path from the start point to the end point for each chromosome, thus resulting in a gapless assembly of the *indica* rice genome (Fig. 1D). Here we found MH63RS2 served as a good reference for locating and orienting contigs to the 12 chromosomes to facilitate assembly. Alternatively, genetic maps (Catchen et al., 2020), HIC (Dudchenko et al., 2017), or other biological methods can also be used to anchor contigs.

**Figure 1.**
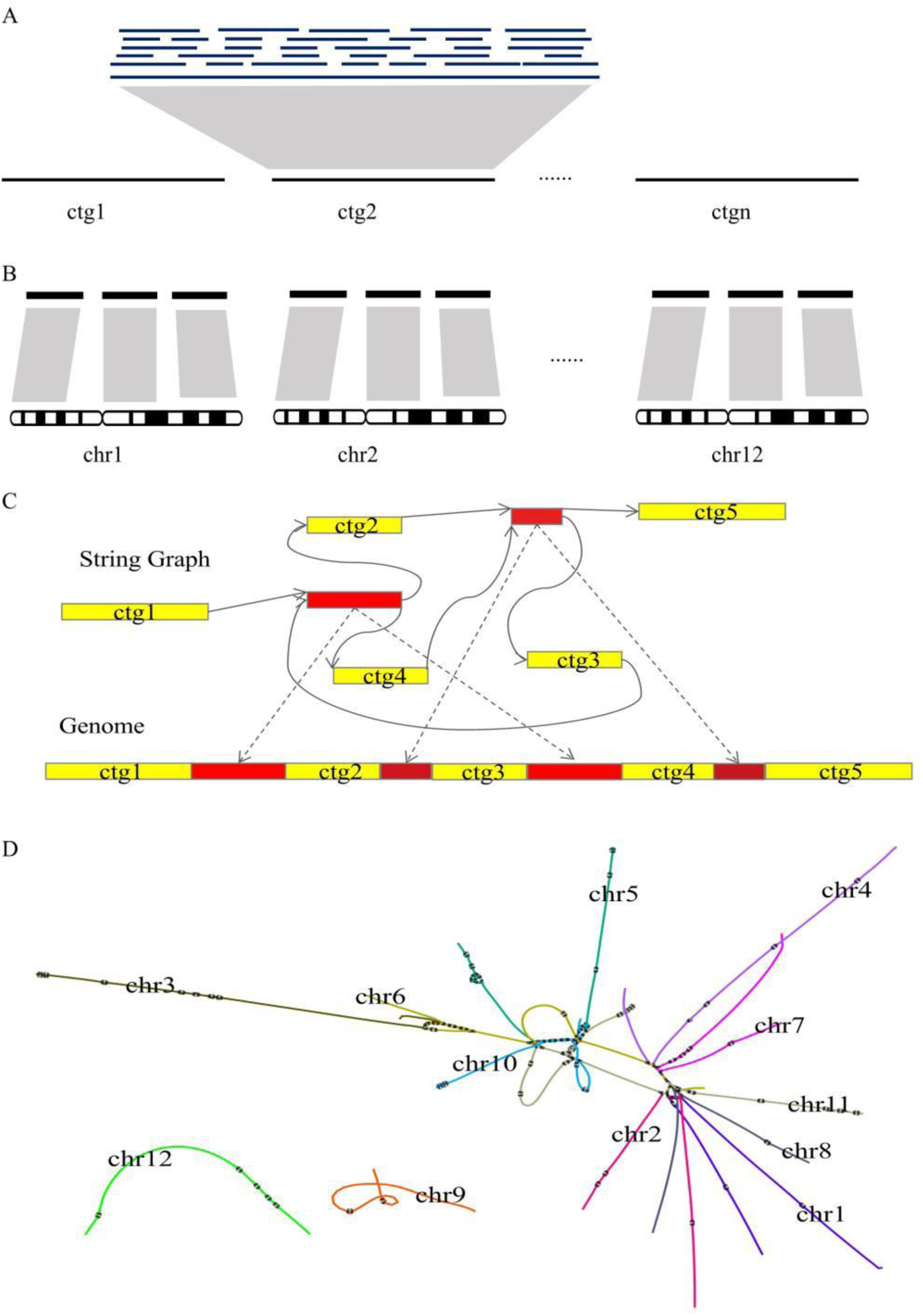
Schematic depiction of the assembly process. (A) Whole-genome assembly from High-fidelity (HiFi) reads. (B) Anchoring WGS contigs onto chromosomes. (C) String graphs were reconstructed using edges (sg_edges_list) and overlap information (pread.m4) generated by pb-assembly. Contigs were linked following their order on each chromosome and a unitig (typically a repeat sequence) for each gap between contigs can be found in the string graph. Red blocks indicate gaps; solid lines indicate contig paths; dashed lines indicate gap locations. (D) The final assembly graph. Each color represents one chromosome.

**Table 1.** Summary of different versions of the rice genome assemblies.

|  | MH63KL1 | MH63RS1<sup>a</sup> | MH63RS2<sup>b</sup> | IRGSP-1.0<sup>c</sup> |
| --- | --- | --- | --- | --- |
| **Total length** | 395,822,112 | 359,937,991 | 387,424,359 | 374,422,835 |
| **Contig number** | 12 | 201 | 26 | 971 |
| **Contig length** | 395,822,112 | 359,918,883 | 387,422,159 | 374,304,580 |
| **Largest contig** | 44,998,099 | 7,413,790 | 39,351,290 | 17,510,758 |
| **Contig N50** | 31,823,770 | 3,032,387 | 27,020,111 | 7,711,345 |
| **Contig L50** | 6 | 40 | 6 | 17 |
| **Length of Ns** | 0 | 19,108 | 2,200 | 118,255 |
| **Rate of Ns** | 0 | 0.0053% | 0.0006% | 0.0316% |

<sup>a,b</sup> Assemblies of the *indica* rice Minghui 63 genome in NCBI.
<sup>c</sup> Assembly of the *japonica* rice Nipponbare genome in NCBI.

To assess the quality of the 24 filled gap areas, we compared the gap region sequences in our new assembly with MH63RS2. Among these gap region sequences, 18 were aligned to the correct positions and showed good collinearity (Fig. 2, Fig. S1, Table S3). One gap region was aligned to a previously known gap area in MH63RS2. For the remaining five gap region sequences, we checked the assembly graph and found that these gap sequences all spanned multiple repeat regions, which may require longer sequencing reads to resolve precisely (Fig. S2). Subsequently, we further used Racon (Vaser et al., 2017) to perform three rounds of correction. The final genome, namely MH63KL1, was assembled into 12 contigs corresponding to 12 chromosomes, resulting in a total length of 395.82 Mb with a contig N50 size of 31.82 Mb, which is both longer and more continuous than presently published rice genomes (Table 1, Table S1).

**Figure 2.**
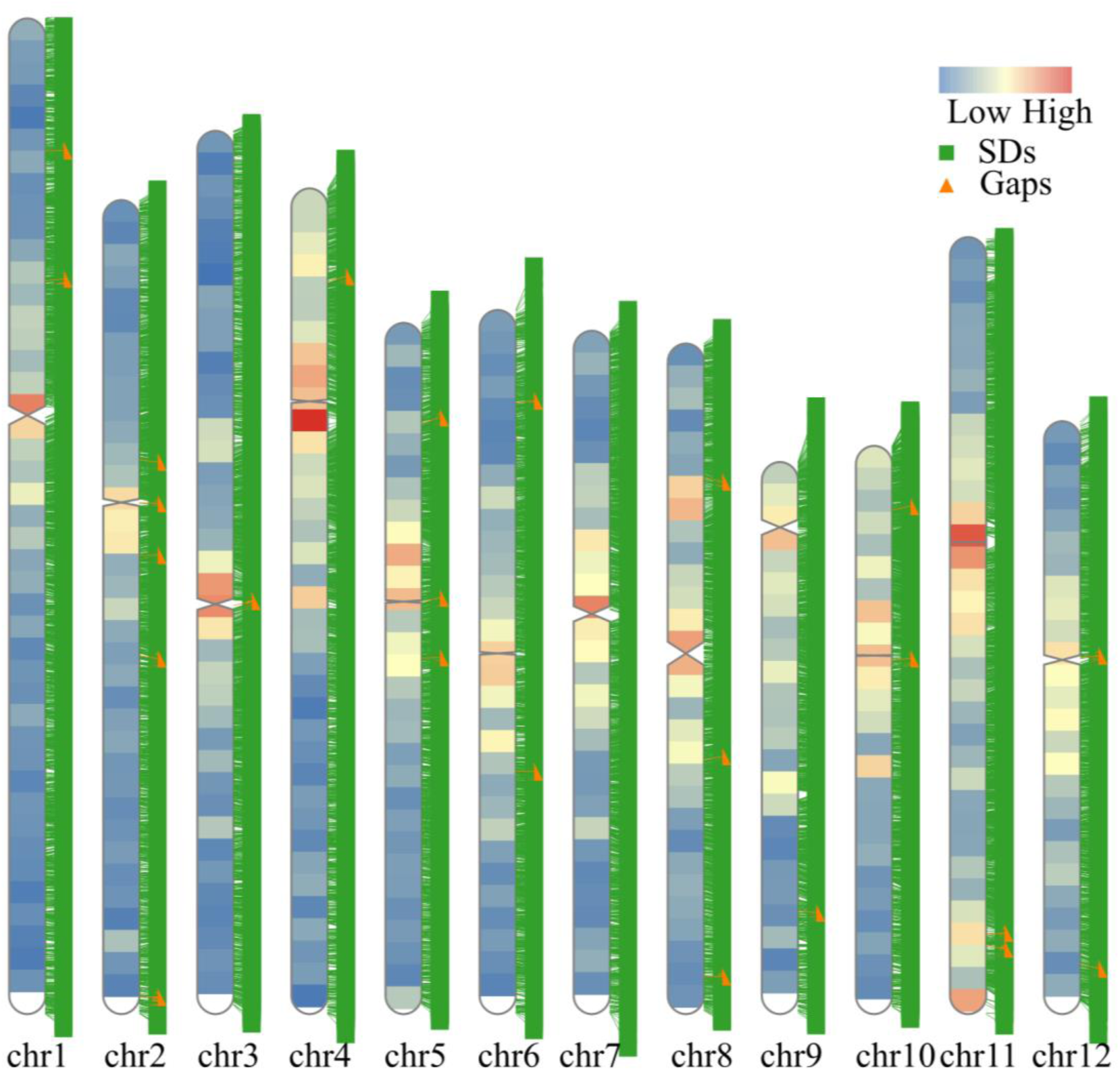
Distribution of repeat sequences and gap positions in MH63KL1. The overlaid heatmap shows the TE density in a 1-Mb window. The tack labels refer to segmental duplications (SDs, green boxes) and the original gap locations (orange triangles).

In order to evaluate the quality of our assembly, Illumina paired-end reads were mapped to our assembly with a mapping rate of 99.34% and a coverage of 98.98% (Table S4). We also mapped 2,045 full-length cDNA sequences to the assembly, among which 1,971 (96.38%) could be mapped with greater than 50% coverage (Table S5). We used NUCmer (Marcais et al., 2018) and MCScan (Tang et al., 2008) to analyze collinearity between our assembly and MH63RS2, and found a high degree of synteny across the whole genome (Fig. S3). The completeness of gene regions of our assembly was evaluated using Benchmarking Universal Single-Copy Orthologs (BUSCO) (Simao et al., 2015). Of the 1614 single-copy orthologs identified in embryophytes, 98.6% were complete in our assembly, which indicates a higher degree of completeness than MH63RS1 (93.1%), MH63RS2 (98.0%), and IRGSP-1.0 (98.3%) (International Rice Genome Sequencing, 2005). The results also indicate that our assembly contains the fewest missing (1.1%) and fragmented (0.3%) single-copy orthologs (Fig. S4, Table S6). LTR-RT annotation of our assembly revealed an LTR assembly index (LAI) score of 22.72, which matches the current gold standard (Fig. S5) (Ou et al., 2018). Finally, we compared our assembly with the sequences of 12 BACs obtained from GenBank, each containing one of the centromeres from rice (http://rice.plantbiology.msu.edu/annotation_pseudo_centromeres.shtml). All the BAC sequences were successfully aligned to our assembly in the correct order and with more than 90% coverage. Since most of the BAC sequences were incomplete, their corresponding areas in our assembly were actually longer (Fig. S6, Table S7). This implies that our assembly of the centromere regions was likely more reliable than sequences of the corresponding BACs.

From our assembly, a total of 36,149 PCGs were predicted, with a mean coding sequence size of 1,146.49 bp and an average of 4.36 exons per gene, similar to the predicted gene sizes in the genomes of other grass species (Table S10, Table S11). Of the PCGs, 99.8% were functionally annotated based on information from the NR (non-redundant, NCBI), SwissProt (Bairoch and Apweiler, 1999), InterPro (Hunter et al., 2012), Pfam (Punta et al., 2012), and KEGG databases (Kanehisa et al., 2014) (Table S12).

We also determined the locations of the centromeres in our assembly by searching for sequences homologous to the highly repetitive 155–165 bp CentO satellite DNA (Cheng et al., 2002). We found that the locations were consistent with previously published results (Zhang et al., 2016) (Table S13). Taken together, these results illustrate the high quality, reliability, and accuracy of our assembly.

### SDs and functional diversification of paralogous gene pairs

SDs are genomic segments >1 Kb that are repeated within the genome with at least 90% sequence identity (Bailey et al., 2001). SDs often carry many duplicated genes and can thus serve as variable hotbeds of gene functional innovation. Due to limitations in the assembly technology, SD regions are often incorrectly assembled, collapsed into a single representative sequence, or completely missed. Consequently, this missing or incorrect information imposes limits on our operability in understanding the structure and evolution of a genome. Our gapless MH63KL1 rice genome provides an opportunity to accurately characterize SDs. To this end, we used SEDEF in this study to analyze the SDs in the rice genome (Bailey et al., 2001), which resulted in the identification of a total of 22,447 SDs, equivalent to 89.68 Mb in length, and accounting for 22.66% of the genome (Fig. 2, 3A, Table S14). We found that SDs are not randomly distributed across the rice genome, but rather, are preferentially located at higher frequencies on chr4, chr10, chr11, and chr12, and at lower frequencies on chr1, chr2, and chr3, suggesting that chr4, chr10, chr11, and chr12 may have a previously unrecognized contribution to the evolution of rice (Table S14).

**Figure 3.**
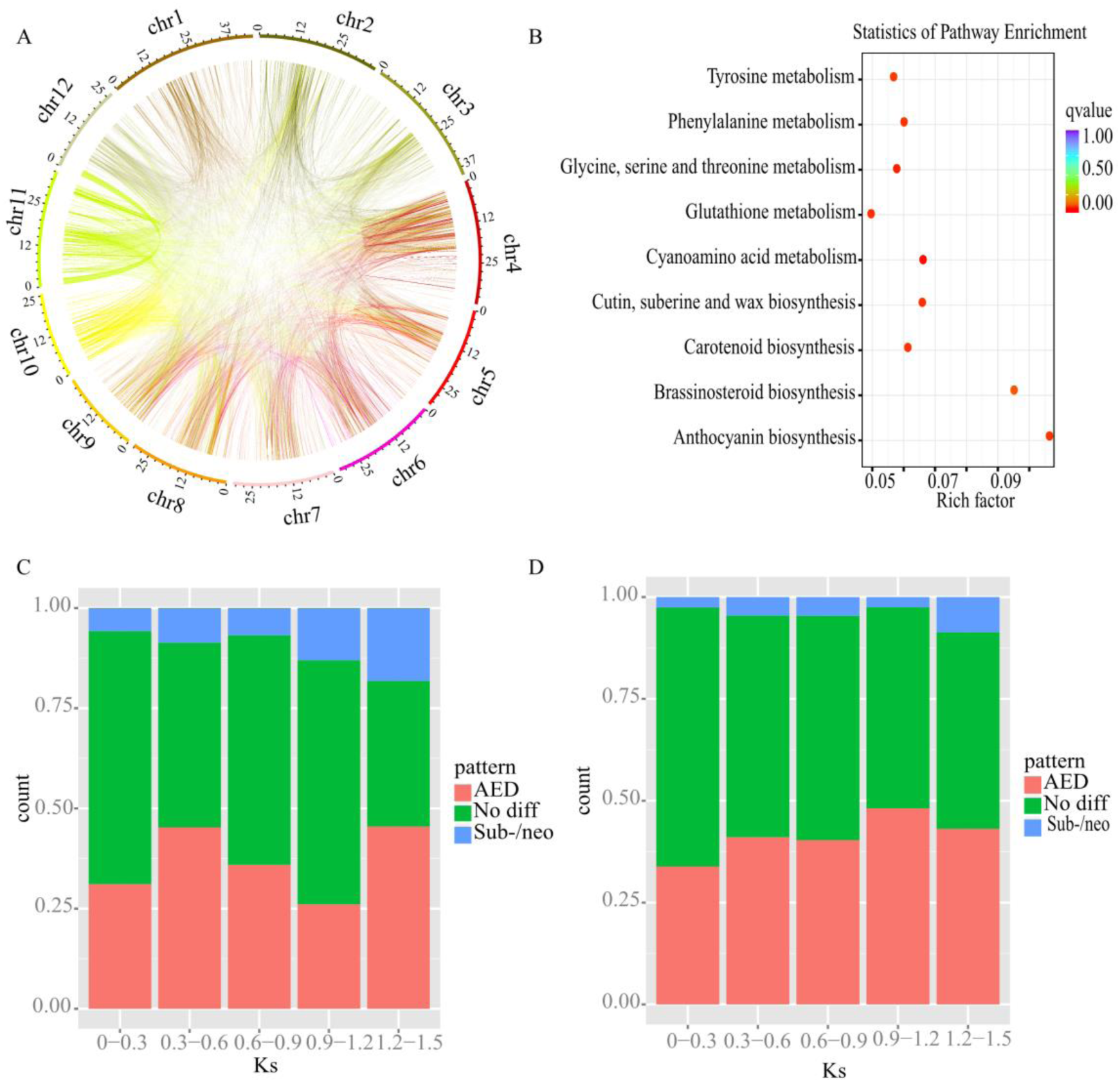
Identification of SDs and their evolutionary impact on the rice genome. (A) Segmental duplicated regions in Minghui 63. Only SDs blocks greater than 5000 bp are shown. (B) Scatterplot of KEGG enrichment for the sub-/neo-functionalized genes in Minghui 63. (C, D) The proportion of each category of duplicated gene pair expression patterns located within 1 Mb in the same chromosomes (C) and on different chromosomes (D) across different Ks values. The X-axis indicates Ks boundaries. The Y-axis is the proportion of duplicates.

We then identified duplicated genes in the SD regions. We initially performed an all-vs-all comparison to find the potential paralogous gene pairs using BLASTP (version 2.2.26) (Altschul et al., 1997) with an E-value threshold of 1e-5. A total of 2,646 pairs were identified in SD regions. For each pair of SD regions harboring paralogous gene pairs, we calculated the Ks values of the gene pairs as proxies for the generation time of their corresponding SDs (Fig. S7). Our result revealed that most of the SDs are relatively recently produced.

The creation of SDs results in a large number of duplicated genes, which can accelerate genome evolution (Conant and Wolfe, 2008). Although most young duplicated genes tend to be functionally redundant and thus are susceptible to loss-of-function mutations that degrade one of the copies into a pseudogene, some duplicated pairs retain their encoded function and continue to express in the long term (Lan and Pritchard, 2016). To determine the fates of these duplicated genes in the SD regions, we analyzed their expression profiles and classified these duplicates into three categories. A gene pair is defined as potentially sub- or neo-functionalized if one gene expresses at a significantly higher level than the other in at least one tested tissue. We refer to those pairs as asymmetrically expressed duplicates (AEDs) when the expression level of one of the pair is higher in at least two tissues and not lower in other tested tissues than its sister gene. The remaining duplicates are classified as having no difference (non-difference) (Lan and Pritchard, 2016).

From the 2,646 gene pairs, 150 were identified as sub-/neo-functional gene pairs, 1,505 as non-difference pairs, and 991 as AEDs (Fig. S8, S9, S10). Then, we analyzed the proportion of each expression pattern category across different Ks values. With increasing time, the proportions of sub-/neo-functional gene pairs and AEDs also gradually increase, while the proportion of no-difference pairs decreases. This finding suggests that, although it is unable to predict which copy of a duplicated gene pair would be preferentially expressed after SD creation, one of them is likely to be subsequently pseudogenized or sub-/neo-functionalized (Fig. S11). Analysis of the positions of duplicated gene pairs showed that more no-difference pairs and sub-/neo-functional pairs are in tandem SDs, and more AEDs are in interspersed SDs (Fig. 3C, 3D). One possible explanation for such a difference is that tandemly repeated genes in the genome may be co-regulated, which facilitates their co-evolution, whereas chromosomally distant duplicated genes may evolve independently.

Kyoto Encyclopedia of Genes and Genomes (KEGG) pathway analyses (FDR <0.05) (Kanehisa et al., 2014) showed that sub-/neo-functionalized genes are mainly enriched in metabolic pathways which may contribute to stress resistance, such as glycine, serine and threonine metabolism (map00260); cutin, suberine and wax biosynthesis (map00073); glutathione metabolism (map00480); anthocyanin biosynthesis (map00942); phenylalanine metabolism (map00360); tyrosine metabolism (map00350); and brassinosteroid metabolism (map00905). Further analysis revealed that the genes encoding 9-*cis*-epoxycarotenoid dioxygenase, glutathione *S*-transferase, primary-amine oxidase, very-long-chain (*3R*)-3-hydroxyacyl-CoA dehydratase, and catalase are duplicated, suggesting that the abundance of functional genes related to the environmental adaptability of rice was expanded by the production of SDs (Fig. 3B, Table S15).

### Active TEs promote rice evolution

We identified a total of 157.16 Mb (39.70%) of TEs in our assembled genome, which was close to the percentage of TEs reported in other published rice genomes (Table S8, Table S9). The predominant TE type is LTR elements, which represents approximately 71.36% (112.13 Mb) of the total length of TEs. We analyzed the genomes of Minghui 63 (MH63KL1) and Nipponbare (IRGSP-1.0) and identified 5,516 and 4,706 LTR-RTs, with a total length of 53.65 Mb and 43.50 Mb, respectively (Table S16). Based on the differences between the LTRs at the ends of each LTR-RT, we calculated the insertion time of the LTR-RTs in Minghui 63 and Nipponbare genomes. Both genomes were found to undergo a dramatic expansion in LTR-RTs approximately 1.5 million years ago (myr). However, within 0.5 myr, Minghui 63 genome began expanding at a significantly higher rate and the expansion of LTR-RTs may still be ongoing (Fig. 4A, 4B). We next classified LTR-RTs according to RT domains and found that the *Del* family in Minghui 63, in particular, exhibits significant expansion (Table S17). Analysis of the divergence between TEs and consensus sequences showed an increase in recent insertion events of TEs such as *En-Spm, MuDR, Tc1, Copia*, and *ERV* (Fig. S12). The above results indicate that TEs in Minghui 63 have a lineage-specific expansion, which may partially explain the larger genome of Minghui 63 compared with Nipponbare. In addition, these findings also suggest that TEs in Minghui 63 are relatively active.

**Figure 4.**
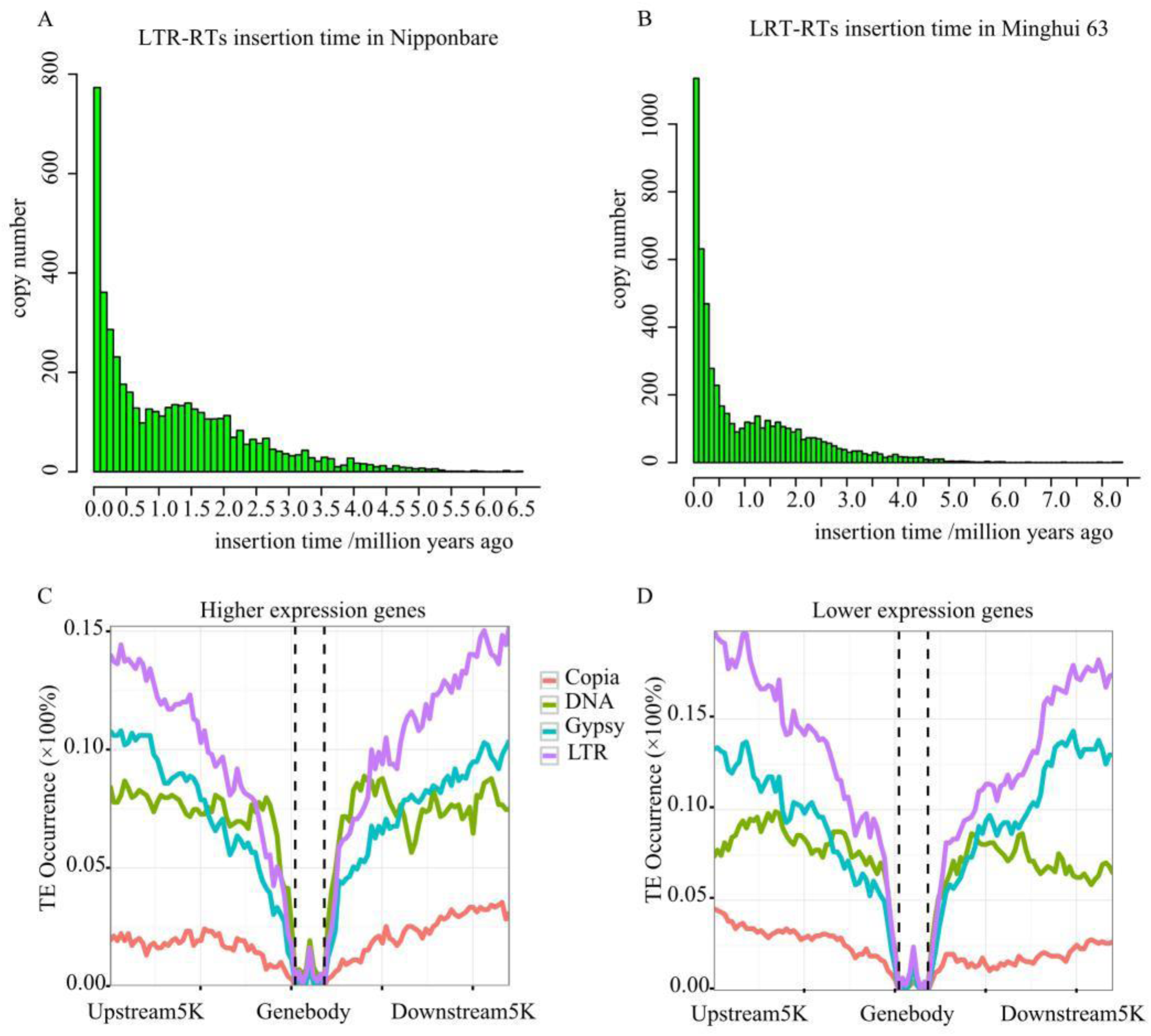
LTR insertions and their effect on gene expression. (A, B) The insertion time of LTR-RTs in Nipponbare (A) and Minghui 63 (B). (C, D) Distributions of Copia, Gypsy, LTR, and DNA transposons across higher (C) and lower (D) expression genes and their flanking regions (±5 kb).

TEs can also substantially contribute to shaping gene expression. TE insertions within the coding regions of a gene can result in loss of function or an aberrant protein. Insertions in introns can change the splicing pattern of transcripts and thus the function of genes. TEs can also affect gene expression by providing novel promoters or regulatory regions. The epigenetic profile of the TEs can alter gene expression as well (Hirsch and Springer, 2017). Given these possibilities, we analyzed the occurrence of TEs around AEDs. We found that, between the duplicated genes, there are higher abundances of all the types of TEs near the copies with lower expression levels (Fig. 4C, 4D).

### SDs can enhance rice disease resistance traits

Plants have evolved a series of defensive mechanisms in the process of resisting pathogen infection, primarily through various disease resistance genes that recognize pathogens and block their invasion (Wan et al., 2012). For example, the nucleotide-binding site with leucine-rich repeat (*NBS-LRR*) genes comprise one of the largest gene families in plants and play a central role in plant resistance to biotic stress (McHale et al., 2006).

Compared with *japonica* rice, *indica* rice has a higher resistance to biotic stresses such as rice blast (Wu et al., 2017). We identified the *NBS-LRR* genes in Minghui 63 and Nipponbare, and found that they are significantly expanded in Minghui 63 compared to Nipponbare (487 versus 349) (Fig. S13, Table S18). We then counted the distribution of *NBS-LRR* genes on each chromosome, and found that they were expanded significantly on chr11 in particular (Table S19). This finding suggests that chr11 may play a major role in disease resistance in rice.

We then identified the orthologous genes between Minghui 63 and Nipponbare and found that 246 *NBS-LRR* genes are shared by Minghui 63 and Nipponbare, 241 are specific to Minghui 63, and 103 are specific to Nipponbare (Fig. S14). We then analyzed the 241 Minghui 63 specific *NBS-LRR* genes, and found that 71 were located within collinear block pairs of SDs. Most of these SDs are tandem repeats except for a few that are located on different chromosomes (Table S20). These results indicate that the tandem SDs likely participate in the expansion of *NBS-LRR* genes and especially in driving evolutionary adaptations to resist pathogen infection in rice. Next, we analyzed the expression profiles of the paralogous genes in the 71 collinear blocks and found that most of the paralogous gene pairs either exhibit no difference in expression or express asymmetrically in different tissues (Fig. S15). This finding is similar to the expression trend we observed for all paralogous genes in SDs in Minghui 63. Furthermore, a large number of *NBS-LRR* genes located in SDs probably suggest a dose effect to successfully resist pathogen infection. A small number of *NBS-LRR* genes may undergo sub-/neo-functionalization and have been co-opted into performing new functions.

Rice blast disease, caused by *Magnaporthe oryzae* (Ascomycota), is one of the most severe and destructive diseases of rice (Valent and Khang, 2010), occurring in almost all rice production regions and commonly reducing yield by 10%-30% (Hossain and Fischer, 1995). We identified the previously cloned and characterized rice blast resistance genes (http://www.ricedata.cn/gene/gene_pi.htm) in Minghui 63 and found that *Pit, Pb1, Piz*, and *Pib* (and their corresponding paralogous genes) located in four collinearity block pairs in SDs. These SDs are all tandem repeats locate on the same chromosome (Table S21). Further analysis revealed that the *Piz* gene had experienced multiple tandem repeat events, two of which occurred in the SD region (Fig. S16, Table S21). Altogether, these findings suggest that the tandem SDs promote the expansion of rice blast resistance genes, which consequently drive new resistance mechanisms against rice blast disease.

### SDs drive evolutionary changes in rice growth and development

Our KEGG pathway analyses (FDR <0.05) of sub-/neo-functionalized genes revealed an enrichment in the zeatin biosynthesis pathway (map00908). Our detailed analysis showed that *cis*-zeatin-*O*-glucosyltransferases (*cZOGTs*) are enriched and sub-functionalized. *cZOGTs*, which catalyzes the *O*-glucosylation of *cis*-zeatin (cZ) and *cis*-zeatin-riboside (Fig. 5A), can result in strong physiological impacts on rice growth and development (Kudo et al., 2012). For example, transgenic rice lines ectopically overexpressing the *cZOGT* genes exhibited short-shoot traits, delayed leaf senescence, and decreased crown root number (Kudo et al., 2012).

**Figure 5.**
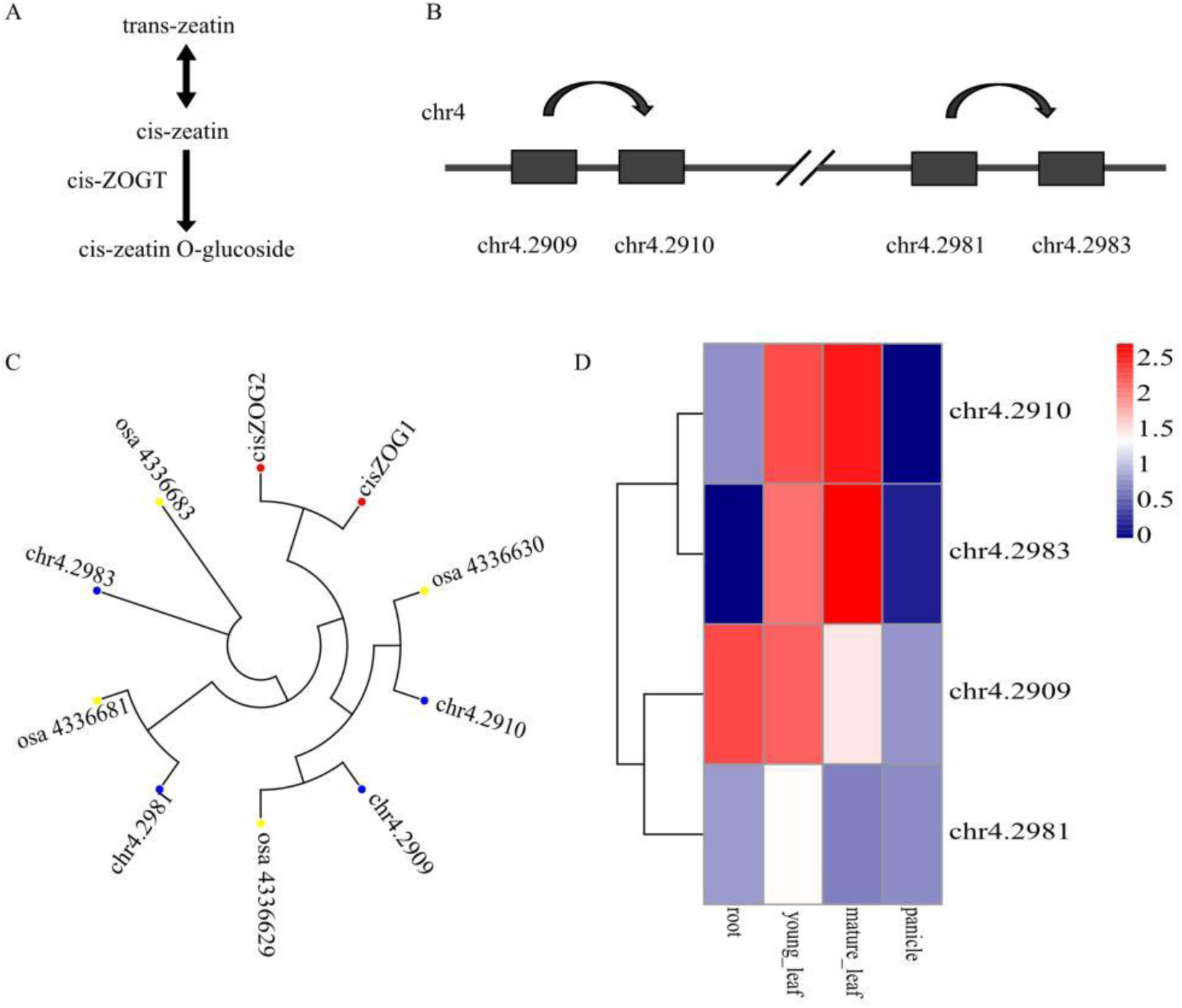
The duplication and diversification of *cZOGTs*. (A) *cZOGTs* catalyze *O*-glucosylation of *cis*-zeatin and *cis*-zeatin-riboside. (B) Schematic depiction *cZOGTs* in collinearity blocks in tandem SDs on chr4. Arrows indicate duplication events. (C) Phylogenetic analysis of *cZOGTs*. Red points indicate corn; blue points indicate Minghui 63; yellow points indicate Nipponbare. (D) Heat map of transcriptional expression for *cZOGTs* in Minghui 63.

We identified *cZOGTs* in Minghui 63 and Nipponbare and found four copies in both varieties, compared with only two copies in corn (Fig. 5B, C). This indicates that *cZOGTs* copy numbers were already increased prior to the divergence of the two varieties of rice. Further analysis revealed that these four genes are distributed on chr4 in two collinearity block pairs of tandem SDs (Table S22), suggesting that *cZOGTs* in rice were doubled by two independent tandem duplication events (Fig. 5B). To further investigate this possibility, we constructed ML trees for *cZOGTs* in Minghui 63 and Nipponbare and found that each group of the tandemly repeated genes are clustered together, which also supports the occurrence of two independent tandem duplication events (Fig. 5C). The gene expression profiles of the *cZOGTs* at different developmental stages showed that two genes are highly expressed in both young and mature leaves, while the third gene is expressed at low levels in all tissues, suggesting its pseudogenization, and the fourth gene is highly expressed in roots and young leaves, suggesting its sub-/neo-functionalization (Fig. 5D). The generation of these genes through segmental duplication strongly suggests that SDs can serve as a source for new genes, some of which may contribute to adaptative or other desirable traits.

## Discussion

Repeat sequences historically represent the most challenging genomic regions to assemble, although they play important roles in the evolution of the genome. With the development of long-read sequencing technology, the continuity of genome assembly has been greatly improved. However, highly accurate, gapless genome assemblies have remained unattainable. In our study, we demonstrated the first strategy to obtain the telomere-to-telomere gapless rice genome, in which the chromosomal position information for contigs is introduced into a string graph. Du et al. (2017) also used chromosomal contig positions to assist assembly, but did not incorporate chromosomal order information of contigs, which is a vital step in our approach. Although their assembly contains only five gaps, there are 14 insertions and deletions (indels) longer than 10 kb, as well as 17 potential indel errors of 1-10 kb (Du et al., 2017). These indels may be introduced by repeat sequences, which cannot be solved by PacBio Continuous long reads (CLRs). Our telomere-to-telomere gapless rice genome is the first gapless genome in higher plants and animals to the best of our knowledge.

Segmental duplications can produce a large number of duplicated genes, thus serving as a source of new genes and subsequently driving rapid adaptative evolution. *Indica* rice has a larger genome than *japonica* rice, likely due to TEs and SDs. The relatively active TEs and newly duplicated genes in Minghui 63 may have an important impact on its agronomic traits. We systematically studied the SDs and TEs in Minghui 63 and identified a total of 89.68 Mb of SDs, accounting for 22.66% of the genome. We also found that SDs in Minghui 63 contain many duplicated genes, a large number of which exhibit no difference in expression, while a small number of which are sub-/neo-functionalized, suggesting that the former might affect plant traits through dose effect and the latter contribute through the gain of new functions.

KEGG enrichment analysis for sub-/neo-functionalized duplicated genes revealed that genes related to disease resistance have expanded, suggesting that the duplicated genes produced by SDs can improve the environmental adaptability of rice (e.g., adaptation to biotic stress or pathogen pressure). TEs can also play an influential role in modulating gene expression. Our analyses showed that TEs are markedly amplified in Minghui 63, and notably, occurred at a significantly higher frequency in the proximity of the genes with relatively lower expression levels in AEDs. We propose that active TEs and SDs work synergistically to promote rice evolution.

Compared with *japonica* rice, *indica* rice has a higher resistance to biotic stresses such as rice blast infection. We found that *NBS-LRR* genes in Minghui 63 have been considerably amplified through duplication, and many of these genes have undergone specialization for pathogen resistance. Notably, 71 *NBS-LRR* genes are located in SDs and most of them are tandem repeat genes. We analyzed the genes related to rice blast resistance and found that four blast resistance genes are located within tandem SDs, suggesting that tandem SDs might drive specific disease resistance adaptations in Minghui 63. In addition to pathogen resistance, we investigated *cZOGTs*, which catalyze *O*-glucosylation of *cis*-zeatin and *cis*-zeatin-riboside, and play important roles in biomass accumulation during the late stage of rice growth and development. Four copies of *cZOGT* genes in rice located on chr4 are found to be derived from two independent tandem repeat events, suggesting that SDs could be potentially engineered through closer study to drive advances in specific desirable agronomic traits.

In our MH63KL1 genome, there are still several overlaps between different chromosomes in the assembly graph. In order to resolve these overlaps, technologies that enable longer sequencing reads but also maintain high fidelity are needed. The development of high-fidelity sequencing technology and the improvement of the genome assembly quality will afford researchers the opportunities to accurately study TEs, SDs, structural variations, and the evolution and expression regulation of duplicated genes.

## Materials and methods

### Genome assembly with PacBio HiFi reads

About 25.3 Gb of Minghui 63 HiFi reads from PacBio Sequel II were downloaded from the NCBI Sequence Read Archive database (SRX6957825). *De novo* assembly of these long reads was performed using pb-assembly (https://github.com/PacificBiosciences/pb-assembly) (Wenger et al., 2019) with parameters ‘-k21 -h850 -e.99 -l2000 -s100 --max-diff 400 --max-cov 400 --min-cov 2 --n-core 24 --min-idt 99.7 --ignore-indels’.

### Gapless assembly

Here we used a new assembly pipeline to obtain a gapless rice genome. The pipeline consists of three steps: (1) anchoring contigs onto chromosomes; (2) linking contig paths to chromosome paths; (3) finding an unitig for each gap in the string graph. (1)

(1) anchoring contigs onto chromosomes

Here we proposed a method of reference-based contig anchoring. NUCmer (Marcais et al., 2018) or MCScan (Tang et al., 2008) was used to map contigs to chromosomes of closely related species (Fig. S3). The positions and orientation of blocks mapped on chromosomes were retrieved. If a contig was matched to multiple chromosomes, we assigned the contig to the chromosome with the largest number of matching blocks. Then contigs were ordered and oriented on each chromosome based on the positions and orientation of their longest blocks on the chromosomes. Then adjacent contigs were linked by 100 Ns. Of course, genetic maps (Catchen et al., 2020), HIC (Dudchenko et al., 2017), or other biological methods can also be used to anchor contigs. Here, the requirement of anchoring accuracy is relatively high and some short contigs can be removed to improve accuracy.

(2) linking contig paths to chromosome paths

First, we extracted the position and orientation of contigs on each chromosome according to the AGP file generated in the previous step to link contig paths generated by pb-assembly (ctg_paths, encoding the graph for each contig and represented by a list of unitigs). Then every chromosome was represented by a list of contig paths and gaps between them. If two contigs were adjacent, they were connected without gaps.

(3) finding an unitig for each gap in the string graph

We reconstructed the string graph using edges (sg_edges_list) and overlap information (pread.m4) generated by pb-assembly. For each gap, we used the starting point of the gap as the center to construct an ego graph and found the paths from the starting point to the ending point of the gap in the ego graph. If there were multiple paths for a gap, we selected the shortest path with the most overlaps. Because there might be assembly errors in the upstream and downstream of gaps, we deleted unitigs near one gap and found the shortest path again if the shortest path for the gap was not found. Finally, if the shortest path was still not found, we kept 100Ns in the corresponding position. The whole assembly graph was shown by Bandage (Wick et al., 2015) (Fig. 1D).

### Assessment of assembly quality

To evaluate the completeness of the assembly and the uniformity of the sequencing, the clean sequencing reads (SRR3234369) of Minghui 63 from short-insert size library were mapped to our assembly using BWA (version 0.7.8) (Li and Durbin, 2009). The mapping rate was 99.96% and the genome coverage was 98.25%.

Benchmarking Universal Single-Copy Orthologs (BUSCO, 4.0.5) (Simao et al., 2015) was used to assess the completeness of gene regions of our assembly using the dataset embryophyta_odb10 (2019-11-20). BUSCO consists of single-copy orthologs derived from OrthoDB in each evolutionary lineage. Of the 1614 single-copy orthologs presented in the embryophytes, 98.6% was complete in MH63KL1, better than the extent in the Nipponbare reference genome (IRGSP-1.0, 98.3%), MH63RS1(93.1%), and MH63RS2(98.0%) (Fig. S4).

In addition to BUSCO,we mapped 2,045 full-length cDNA sequences form Oryza rufipogon W1943 (http://server.ncgr.ac.cn/ricd/dym/ftp.php) to our assembly using BLAT (Kent, 2002). 1,971 (96.38%) of these could be mapped with greater than 50% coverage.

LTR Assembly Index (LAI) uses LTR-RTs to evaluate assembly continuity.The LAI were calculated using LTR_retriever with default parameters (Ou et al., 2018).

Finally,12 BAC sequences of centromere regions obtained from the NCBI GenBank and 24 gap region sequences were aligned to our assembly using LASTZ(http://www.bx.psu.edu/~rsharris/lastz/). The detailed information of sequence alignments was shown in Fig. S1, Fig. S6, Table S3, and Table S7.

### Collinearity analysis

MCScan (Tang et al., 2008) was used to identify the collinearity of genes between MH63KL1 and MH63RS2 with parameters (--cscore=0.99) (Fig. S3B). NUCmer was used to align MH63KL1 and MH63RS2 with parameters (--mum, --mincluster=200, - -minmatch=100). Delta-filter was then used to get the 1-to-1 alignments with parameters (-i 95, -l 100, -1). The dot plot was showed using mummerplot (Fig. S3A) (Marcais et al., 2018).

### Repeats annotation

Transposable elements of Minghui 63 genome were annotated through a combination of *ab initio* prediction and homology searching. We first used LTR_FINDER (Xu and Wang, 2007), RepeatScout (Price et al., 2005), and RepeatModeler to build a consensus sequence library. Repeatmasker (http://www.repeatmasker.org/RepeatModeler.html) was used to annotate the repeat regions based on this library as well as the known Rice TE library (RiTE database, https://www.genome.arizona.edu/cgi-bin/rite/index.cgi) (Copetti et al., 2015). RepeatProteinMask was then used to identify repeat sequences at the protein level. Tandem Repeats Finder (TRF) (Benson, 1999) was used to find Tandem repeats. The divergences of TEs were retrieved from the GFF file produced by RepeatMasker.

### Gene annotation

Protein-coding genes were conducted using a combination of homology-based prediction, *de novo* prediction, and transcriptome-based prediction methods. Proteins sequences from seven plant genomes (*Oryza sativa* L. *japonica*, *Oryza sativa* L. *indica*, *Brachypodium distachyon*, *Setaria italica*, *Zea mays*, *Arabidopsis thaliana*, and *Sorghum bicolor*) were downloaded from Ensemble (release-47). They were aligned to the Minghui 63 genome using the TBLASTN program (Altschul et al., 1990) with an E-value cutoff of 1e-5. The BLAST hits were conjoined using Solar software (Yu et al., 2006). GeneWise (Birney et al., 2004) was then utilized to predict the exact gene structure of the corresponding genomic regions for each BLAST hit. FL-cDNAs and ESTs (RICD,http://www.ncgr.ac.cn/ricd) of *Oryza sativa* L. *indica* and FL-cDNAs of *Oryza sativa* L. *japonica* (Rice Full-Length c et al., 2003) were directly mapped to the Minghui 63 genome and assembled by PASA (Campbell et al., 2006). Gene models created by PASA were used to train the *ab initio* gene prediction programs. Augustus (version 2.5.5) (Urnov et al., 2005), Genscan (version1.0) (Burge and Karlin, 1997), GlimmerHMM (version 3.0.1) (Majoros et al., 2004), Geneid (Blanco et al., 2007), and SNAP (Korf, 2004) were used for *ab initio* gene predictions in the repeat-masked genome. RNA-Seq reads downloaded from the NCBI Sequence Read Archive database (SRR10751892-SRR10751899) were mapped to the assembly using Tophat (version 2.0.8) (Trapnell et al., 2009) with default parameters. Cufflinks (version 2.1.1) (Trapnell et al., 2010) was then used to assemble the transcripts into gene models. All evidence of gene models were combined by Evidence Modeler (EVM) (Haas et al., 2008) into a non-redundant set of gene structures. UTR regions were then identified by integrating transcript evidence using PASA. Finally, gene models were filtered by removing the genes having 20% of their CDS sharing overlaps with TEs and coding region lengths less than 150 bp. The function of protein-coding genes was achieved by searching their protein sequences against the SwissProt (http://web.expasy.org/docs/swiss-prot/guideline.html) and NR databases using BLASTP (Altschul et al., 1997) (E-value 1e-05). Protein domains were identified by searching against the InterPro (http://www.ebi.ac.uk/interpro/, V32.0) and Pfam databases (http://pfam.xfam.org/, V27.0), using InterProScan (V4.8) and HMMER (Finn et al., 2011), respectively. The Gene Ontology (GO, http://www.geneontology.org/page/go-database) terms for each gene was obtained from the corresponding InterPro or Pfam entry. KEGG metabolic pathways were further assigned by BLAST (E-value 1e-05) against the KEGG databases (release 53) (Kanehisa et al., 2014).

### Centromeres identification

HMMER (version 3.1b1) (Finn et al., 2011) was used to identify centromeres. Firstly, we built a hmm file through hmmbuild using a file of 155-165 bp CentO satellite DNA sequences in rice. Hmmsearch was then used to search the location of centromeres of our assembly.

### SDs detection

Briefly, our genome assembly was first soft-masked converting all common and tandem repeats to lower-case letters. SEDEF was then used to detect SDs with the default parameters (Numanagic et al., 2018).

### Ks distribution of the duplicated gene pairs

The protein sequences of the duplicated pairs in SDs were retrieved, and aligned using the MUSCLE program (Edgar, 2004). Subsequently, the protein sequences alignments were converted into CDS alignments based on the coding sequences. We then calculated the numbers of synonymous substitutions per synonymous site (Ks) for these gene pairs based on the MYN method implemented in the PAML program (version 4.9b) (Yang, 2007). The Ks distribution was plotted and displayed using R (version 3.0.1).

### Classification of expression patterns of sister duplicates

RNA-Seq reads of roots, young leaves, mature leaves, and panicles from the NCBI Sequence Read Archive database were mapped to MH63KL1 using TopHat (version 2.0.8) (Trapnell et al., 2009). HTSeq-count was used to calculate the numbers of aligned reads (read counts) for each gene (Anders et al., 2015). Differential gene expression (DEG) analyses among the duplicated gene pairs for each tissue were performed using DESeq2 with FDR cut-off of 0.05 and log2 fold change (FC) cut-off of 1 (Love et al., 2014). We divided the duplicated pairs into three classes according to their expression in four tissues: 1) Sub- or neo-functionalized pairs in which each of the two duplicates was significantly more highly expressed than the other in at least one tissue; 2) Asymmetrically Expressed Duplicates (AEDs) in which one duplicate was expressed higher than its sister in at least two tissues and its expression was not lower than its partner in other tested tissues; 3) The remaining duplicates were classified as no difference pairs (Lan and Pritchard, 2016).

### KEGG enrichment analysis

KOBAS software was used to test the statistical enrichment of differential expression genes in KEGG Pathways with the cutoff set at P < 0.05 (Xie et al., 2011).

### Identification and classification of LTR retrotransposons

We performed *de novo* searches for LTR retrotransposons (LTR-RTs) against our assembly using LTRharvest (Ellinghaus et al., 2008) and LTRfinder. Profile HMM models for domain detection were downloaded from the Gypsy Database (GyDB) (Llorens et al., 2011). LTRdigest (Steinbiss et al., 2009) was used for structure annotation (e.g., PBS, PPT, protein) of LTR-RTs (Liu et al., 2020b; Zhang et al., 2020). The RT domains were used to classify LTR-RT. The nucleotide substitution (λ) between the 5′ and 3′ terminals of intact LTR-RTs were estimated by MUSCLE (Edgar, 2004). The genetic distances (K) were calculated by K = −0.75 ln(1 - 4λ/3). Finally, the insertion time of LTR-RTs was calculated based on the following formula: T = K/2r (r = 1.3 * 10-8 per site and per year) (Ma and Bennetzen, 2004).

### Identification of *NBS-LRR* genes

For each species, the protein sequences were retrieved and searched against the raw Hidden Markov Model (HMM) of NB-ARC family (PF00931) using HMMER (version 3.1b1) (Finn et al., 2011) with default parameters. The high-quality NBS protein sets (< 1E-60) from these initial searches were then self-aligned using mafft (version 7.305) (Katoh and Standley, 2013). The NBS HMMs were then construct using the program of hmmbuild implemented in HMMER (Finn et al., 2011) and were used to search NBS-encoding proteins (Xia et al., 2017). Finally, we identified 487 and 349 NBS-encoding proteins in Minghui 63 and Nipponbare, respectively. PfamScan was used to search these proteins against Pfam-A database (Pfam31.0) (Punta et al., 2012) to identify domains. CC domains were identified using ncoils (Lupas et al., 1991) with the default parameters.

### Identification and evolutionary analysis of rice blast resistance genes and *cis-*zeatin-*O*-glucosyltransferases (cZOGTs) genes

Based on the previously cloned amino acid sequences(http://www.ricedata.cn/gene/), we identified rice blast resistance and *cZOGTs* genes (Kudo et al., 2012) using the BLASTP program (version 2.2.26) (Altschul et al., 1997) with an E-value of ≤1E-10. Only those hits with ≥ 30% identity and ≥ 50% coverage of both query and target were retained.The conserved domains were further examined in the retained sequences. The protein sequences for each gene were individually retrieved and aligned using MUSCLE (version 3.8.31) (Edgar, 2004). Maximum-Likelihood (ML) trees were constructed using RAxML(version 8.0.19) (Stamatakis, 2006) with the PROTGAMMAAUTO model and 100 bootstrap replicates for each alignment. We further checked each evolutionary tree to further filter the identified genes. RNA-seq reads were mapped to our assembly using TopHat (version 2.0.8) (Trapnell et al., 2009). Cuffdiff(version 2.2.1) (Trapnell et al., 2013) was used to compute the FPKM (Fragments Per Kilobase Of Exon Per Million Fragments Mapped) values for each gene. We visualized the gene expression levels using pheatmap (https://cran.r-project.org/web/packages/pheatmap/index.html) for the identified genes.

## Availability of data and materials

The genome assemblies have been deposited in GenBank under accession number CP059874-CP059885. The code used in this study is available at the GitHub repository https://github.com/likui345/PGA.

## Author Contributions

KL, PFL and SL designed the project. KL, WJ, YH, and MK worked on data analyses and developed the gapless assembly pipeline. KL, LZG, PFL and SL wrote the manuscript. All authors read and approved the final manuscript.

## Acknowledgments

We thank Evan E. Eichler (Univ. Washington) for his help with SD analysis.

## Declaration of interets

The authors declare that they have no competing interests.

## SUPPLEMENTARY FIGURES

**Supplemental Figure 1.**
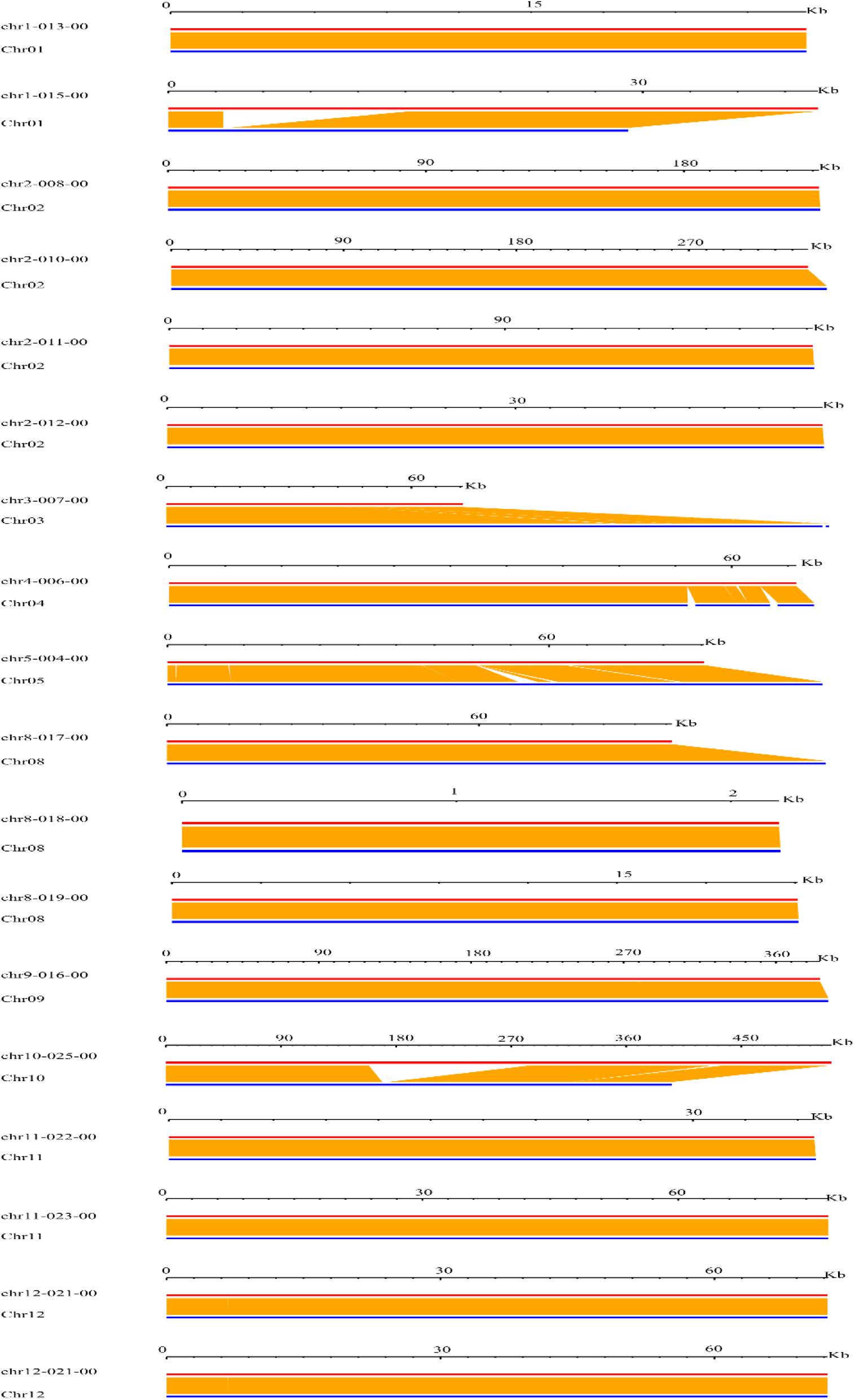
Alignments between 18 gaps of Minghui 63 and MH63RS2. The red line indicates each gap sequence and the blue line represents our assembly. The blocks in white indicate the unfilled gaps. The green blocks show that gap sequences are aligned to the reverse complementary sequence of our assembly.

**Supplemental Figure 2.**
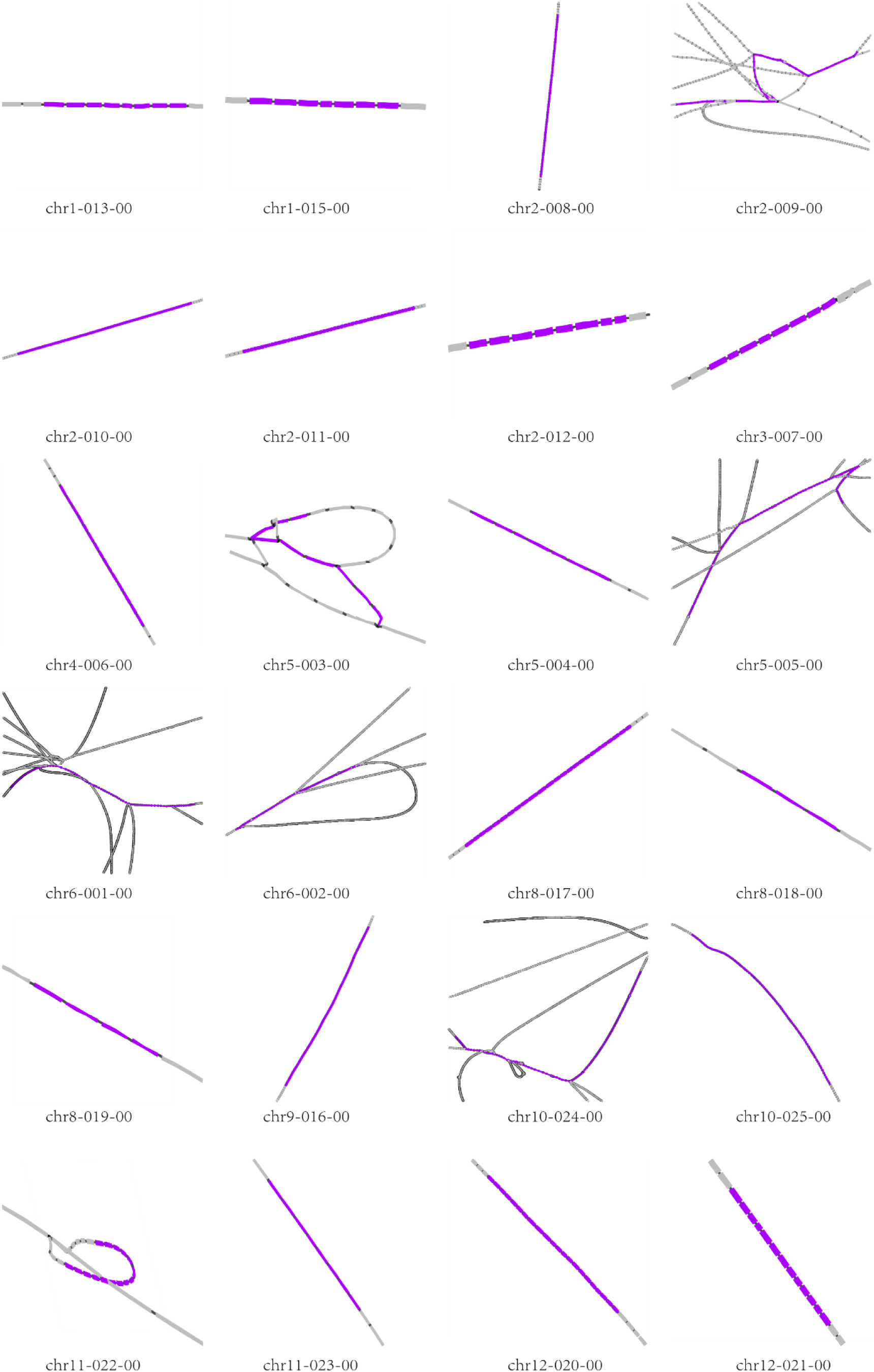
Assembly graphs around 24 gaps. The gaps were marked in purple.

**Supplemental Figure 3.**
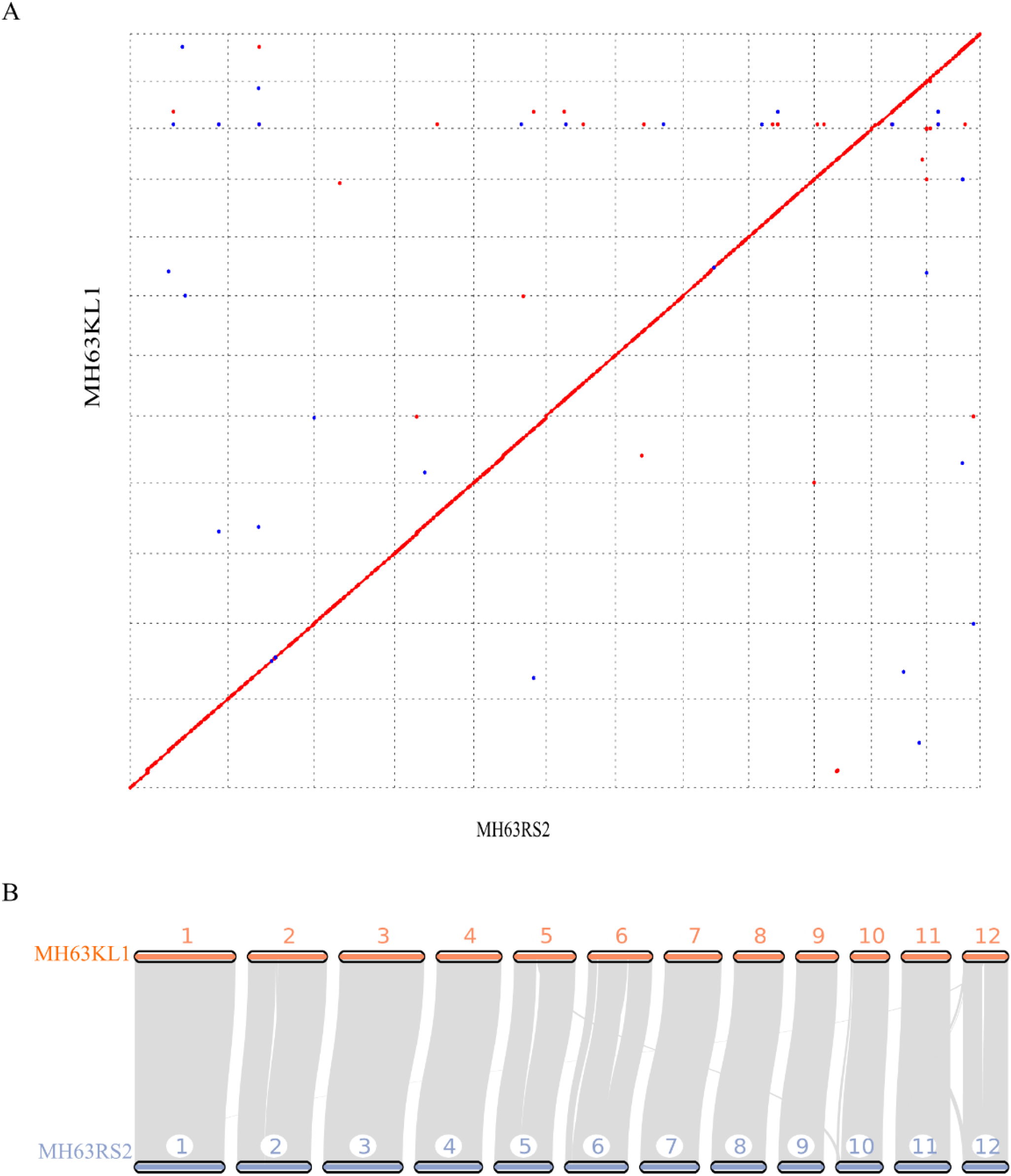
The synteny between Minghui 63 and MH63RS2 across the whole genome. (A) Dot plots of Minghui 63 versus MH63RS2 detected by NUCmer. (B) Syntenic regions between Minghui 63 and MH63RS2 detected by MCScanx.

**Supplemental Figure 4.**
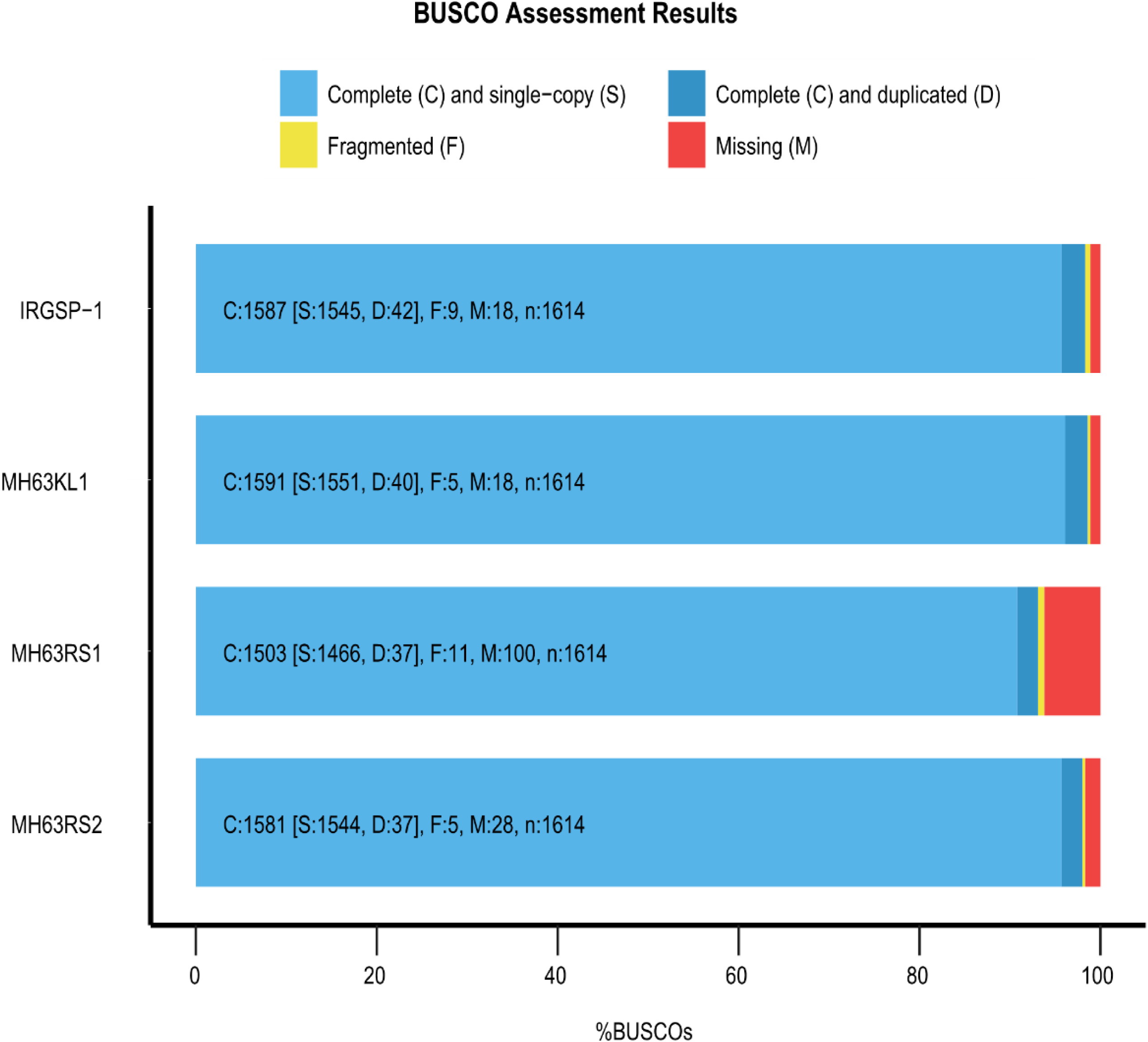
Assessment of the gene coverage rate using BUSCO.

**Supplemental Figure 5.**
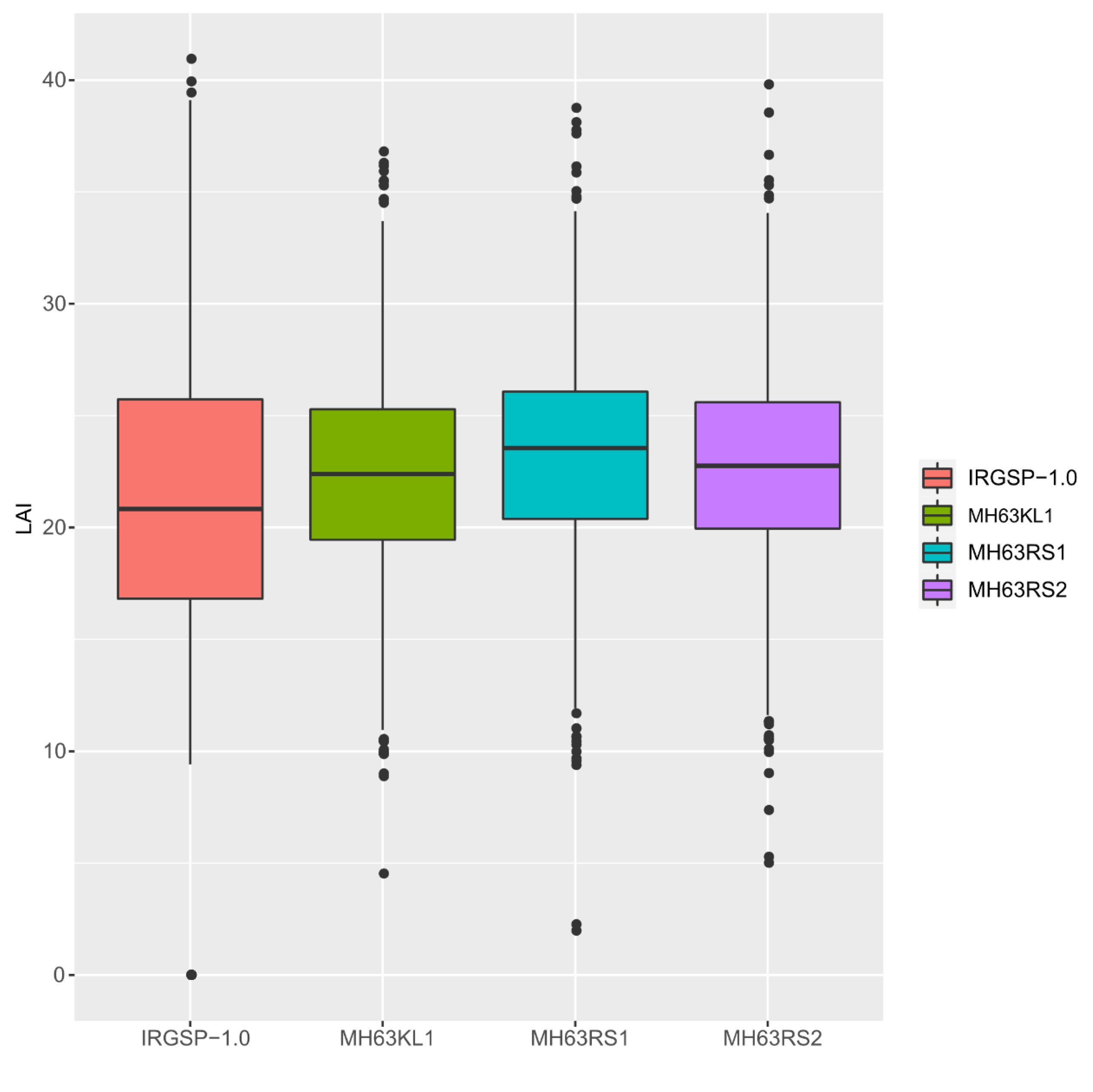
Comparison of LAI scores among four genome assemblies.

**Supplemental Figure 6.**
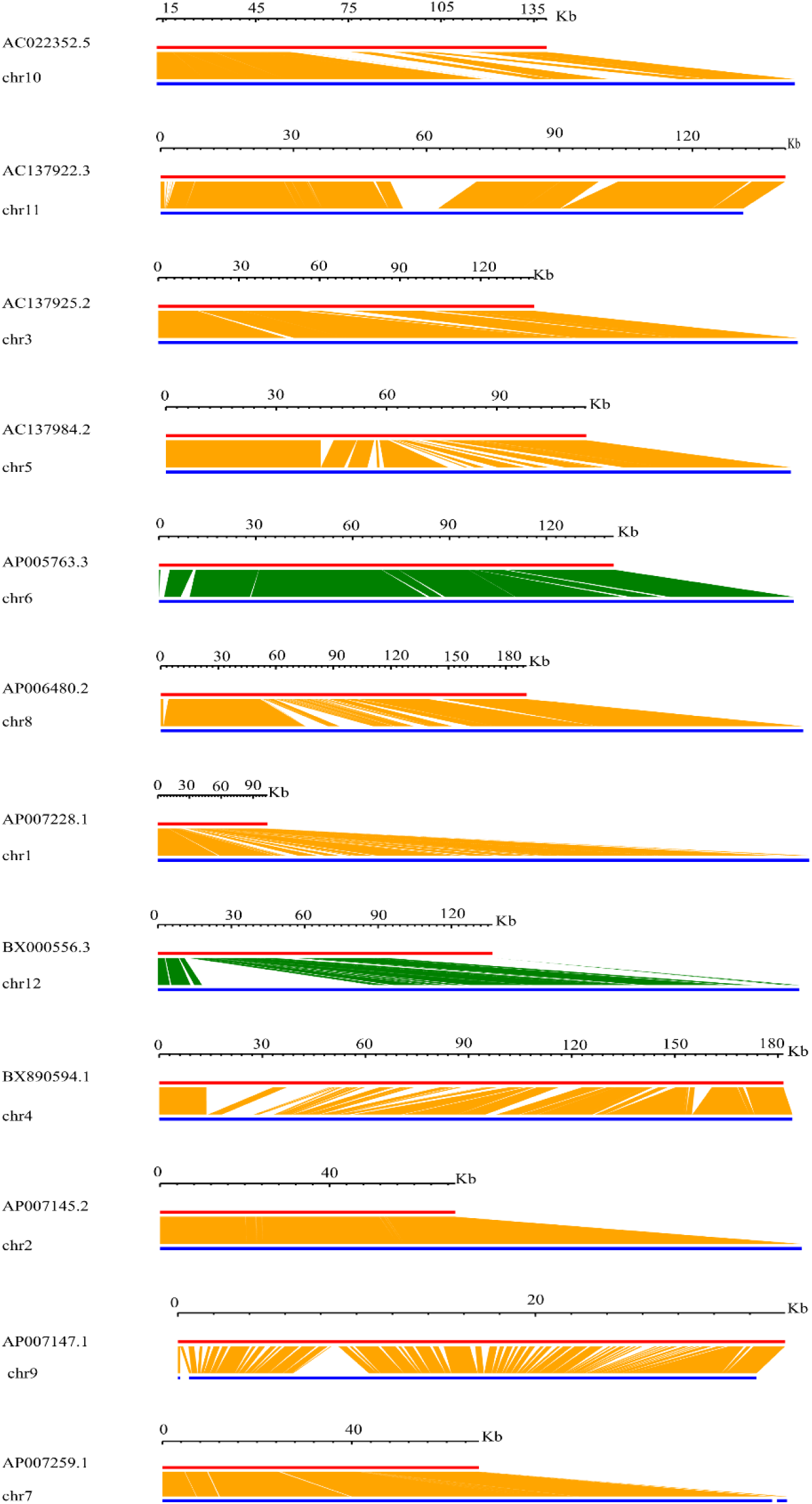
Assessment of the Minghui 63 genome assembly by 12 BACs. The red line indicates each BAC sequence and the blue line represents our assembly. The blocks in white indicate the unfilled gaps. The green blocks show that BAC sequences are aligned to the reverse complementary sequence of our assembly.

**Supplemental Figure 7.**
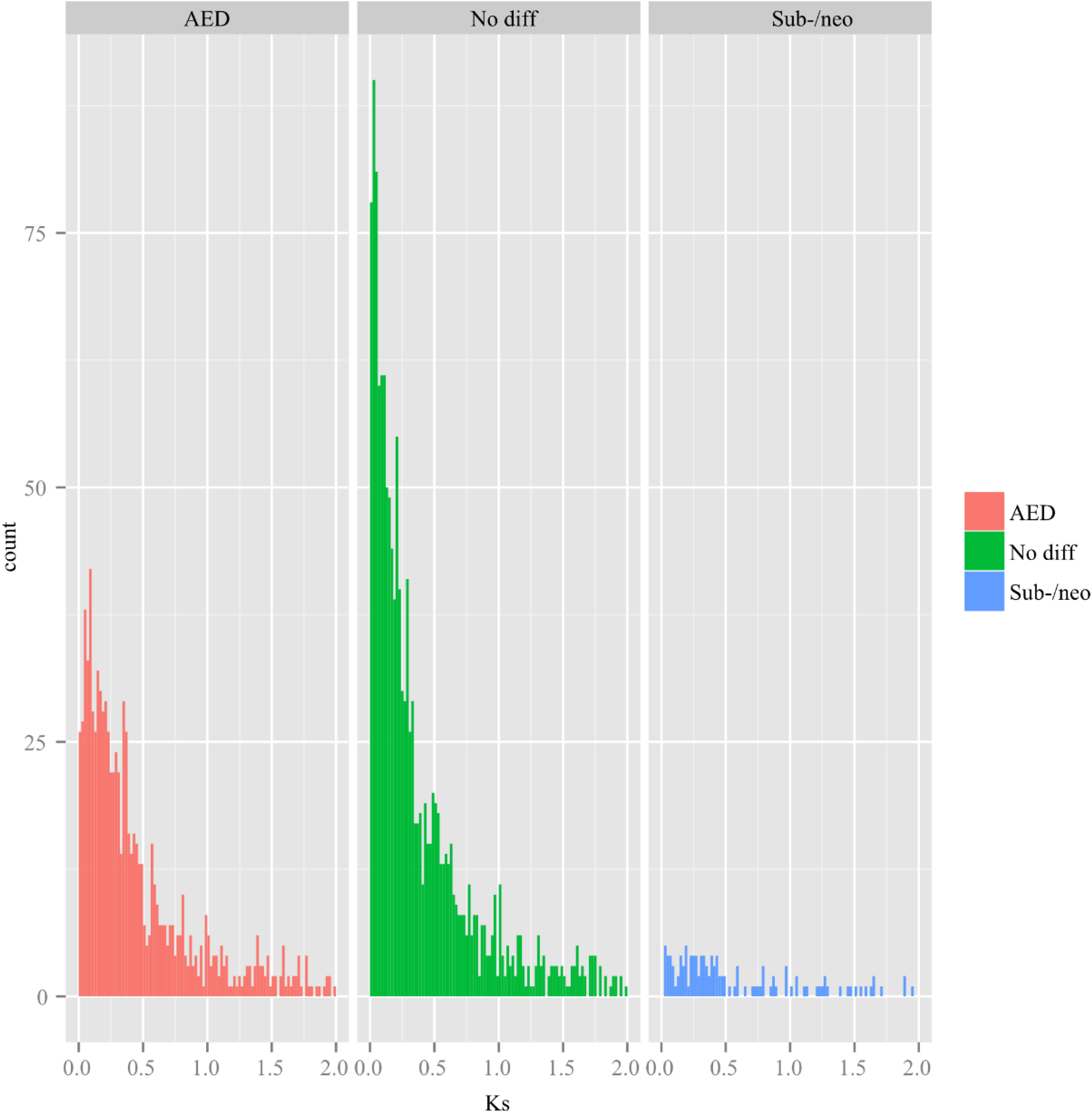
Numbers of different type of duplicate gene pairs across different values of Ks.

**Supplemental Figure 8.**
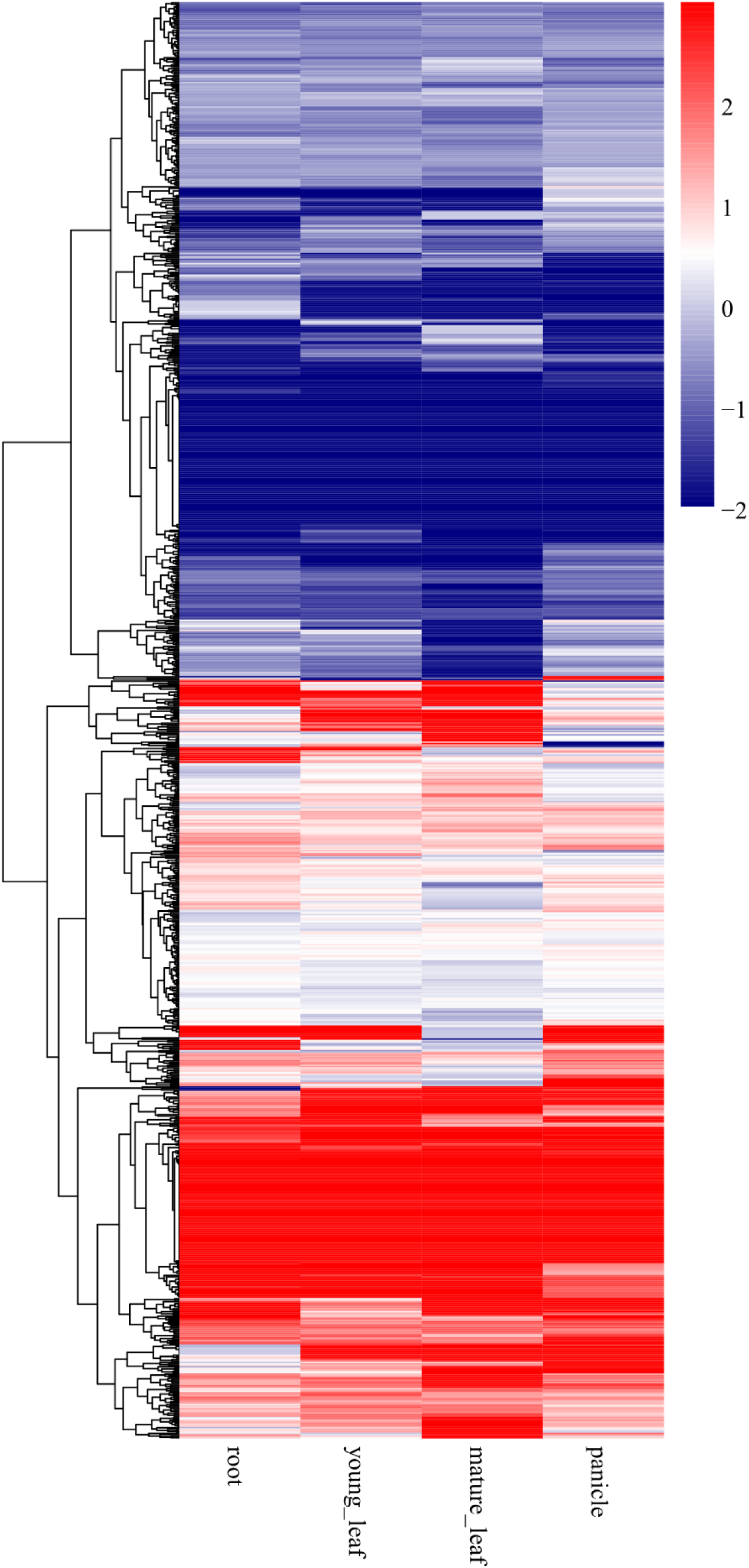
Heat map of expression ratios for AEDs. For each duplicate pair, the ratios show the tissue-specific expression level of each gene relative to its duplicate.

**Supplemental Figure 9.**
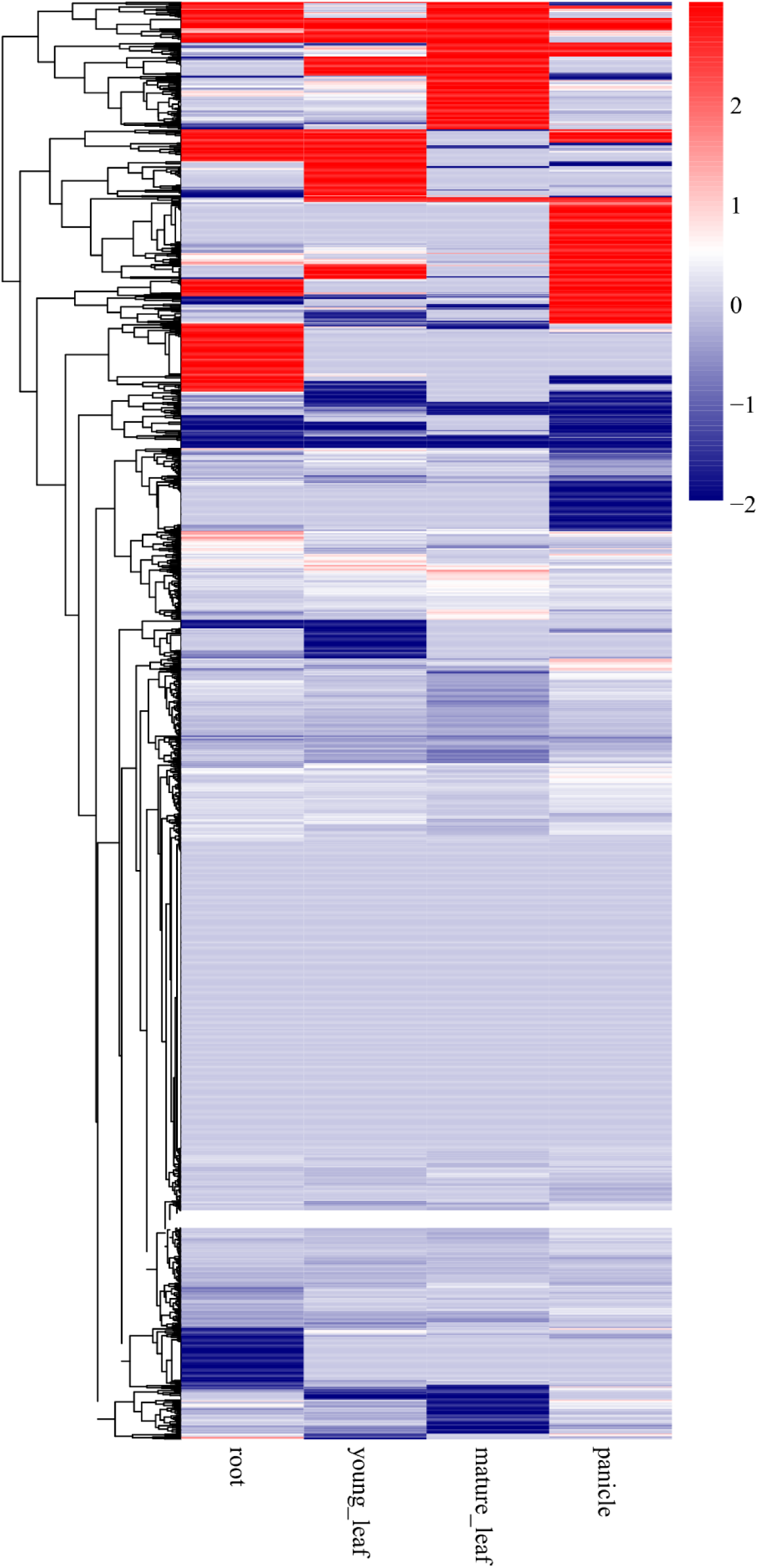
Heat map of expression ratios for no significant difference duplicate pairs. For each duplicate pair, the ratios show the tissue-specific expression level of each gene relative to its duplicate.

**Supplemental Figure 10.**
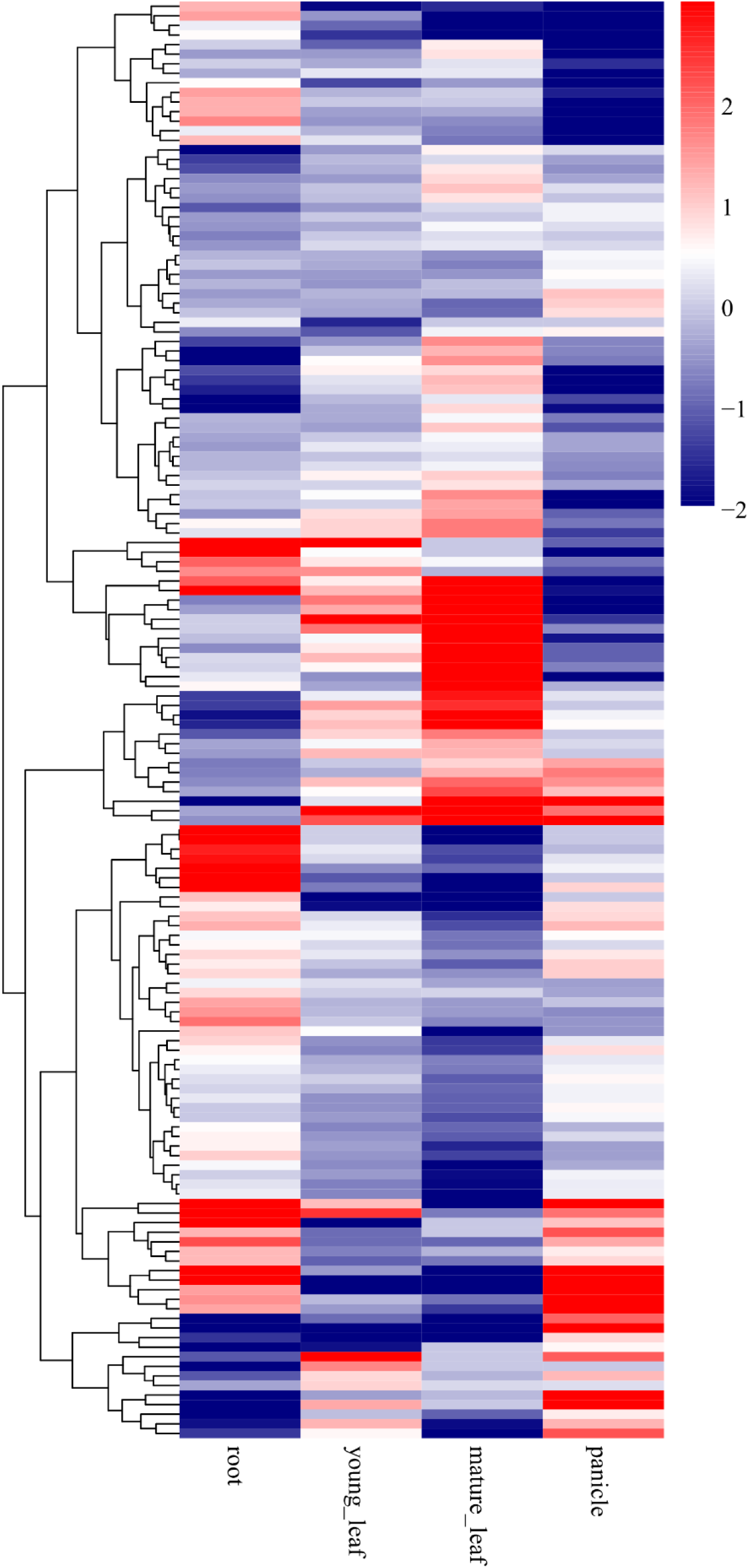
Heat map of expression ratios for subfunctionalization duplicate pairs. For each duplicate pair, the ratios show the tissue-specific expression level of each gene relative to its duplicate.

**Supplemental Figure 11.**
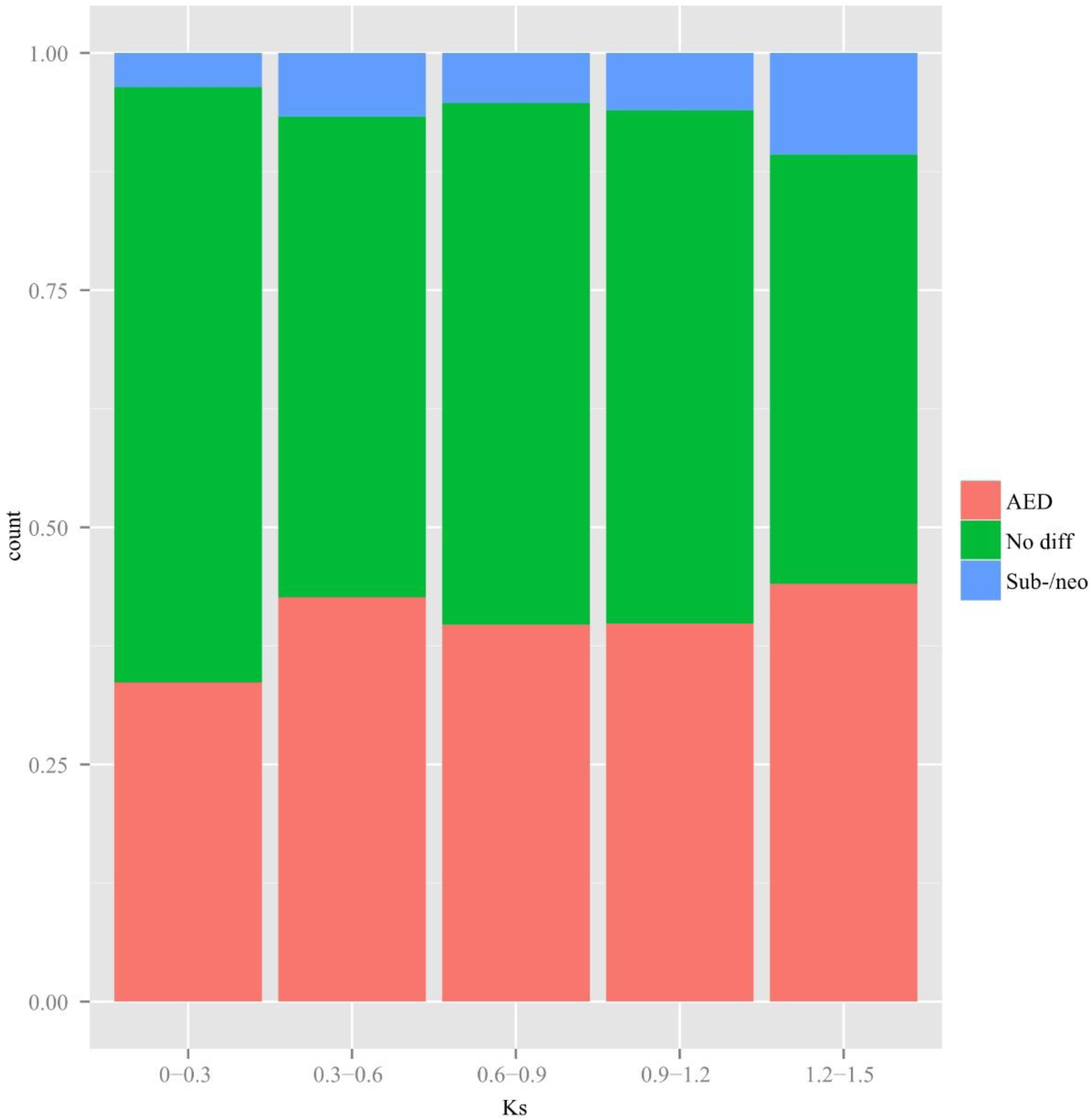
Expression patterns of all the duplicate gene pairs in SDs across different values of Ks.

**Supplemental Figure 12.**
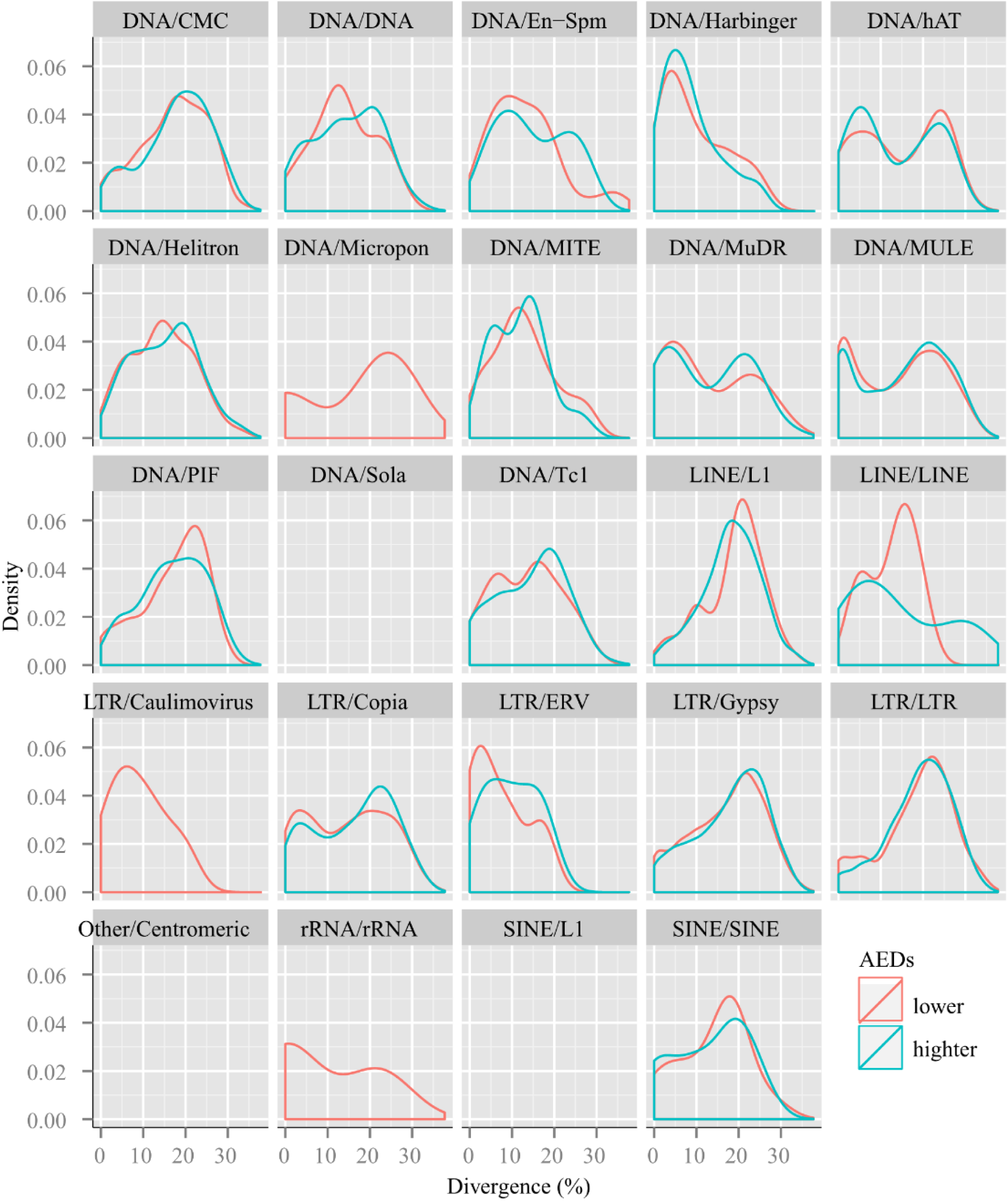
Divergence distribution of classified families of TEs around AEDs. Red lines indicate divergence distribution of TEs around lower expression genes. Green lines indicate divergence distribution of TEs around higher expression genes.

**Supplemental Figure 13.**
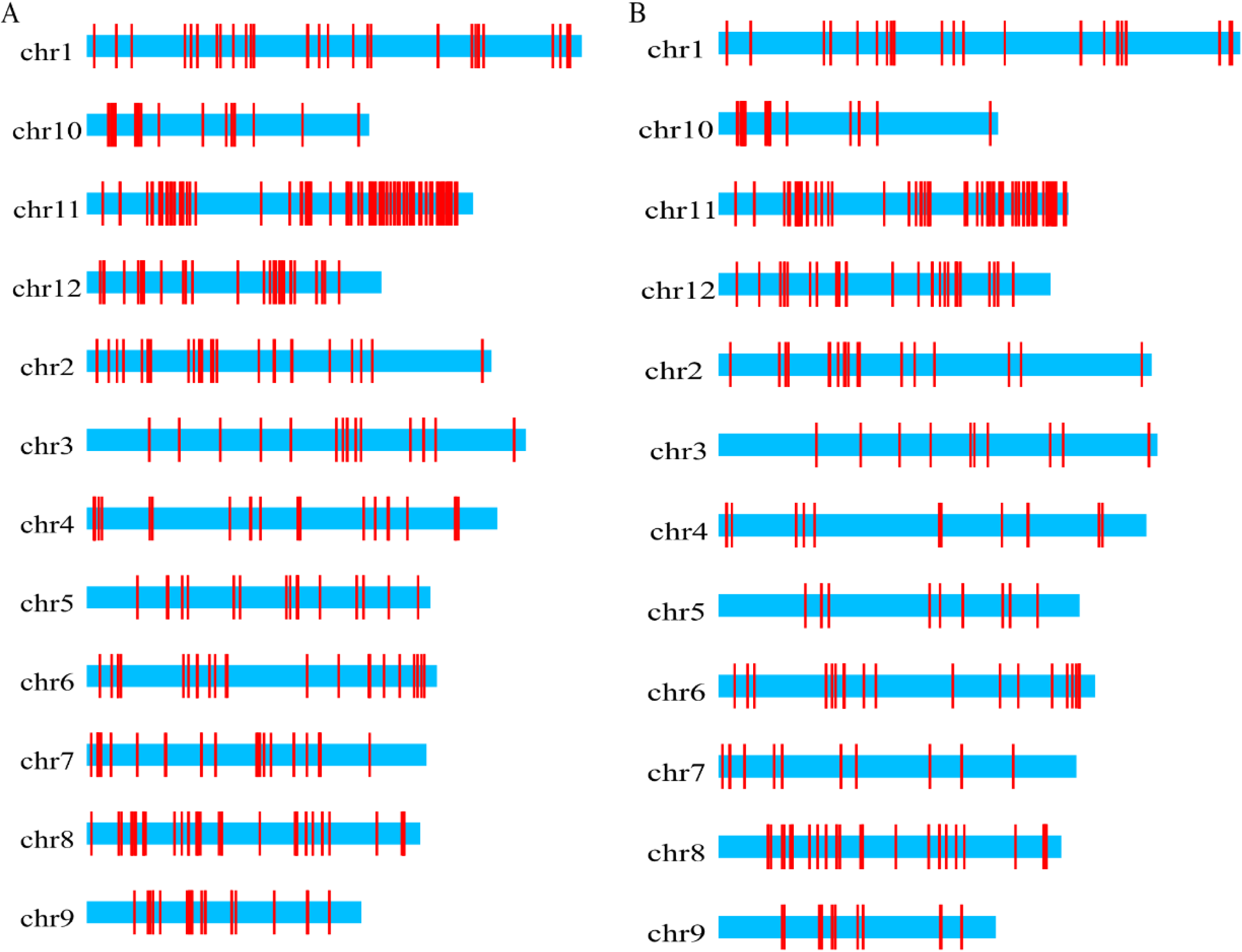
Distribution of *NBS-LRR* genes. A. Distribution of *NBS-LRR* genes in Minghui 63. B. Distribution of *NBS-LRR* genes in Nipponbare.

**Supplemental Figure 14.**
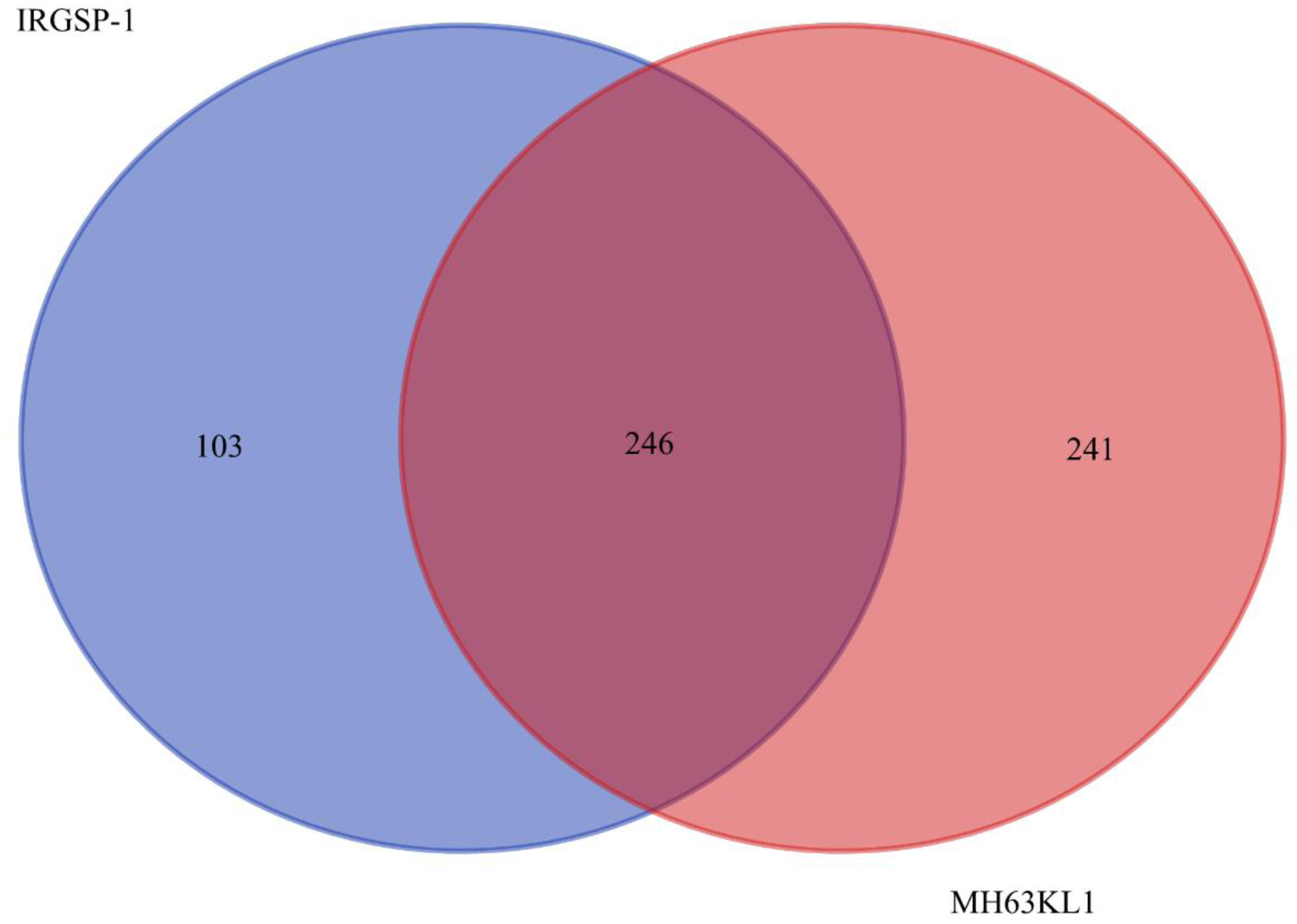
The collinearity of *NBS-LRR* genes in genomes of Minghui63 (MH63KL1) and Nipponbare (IRGSP-1) rice.

**Supplemental Figure 15.**
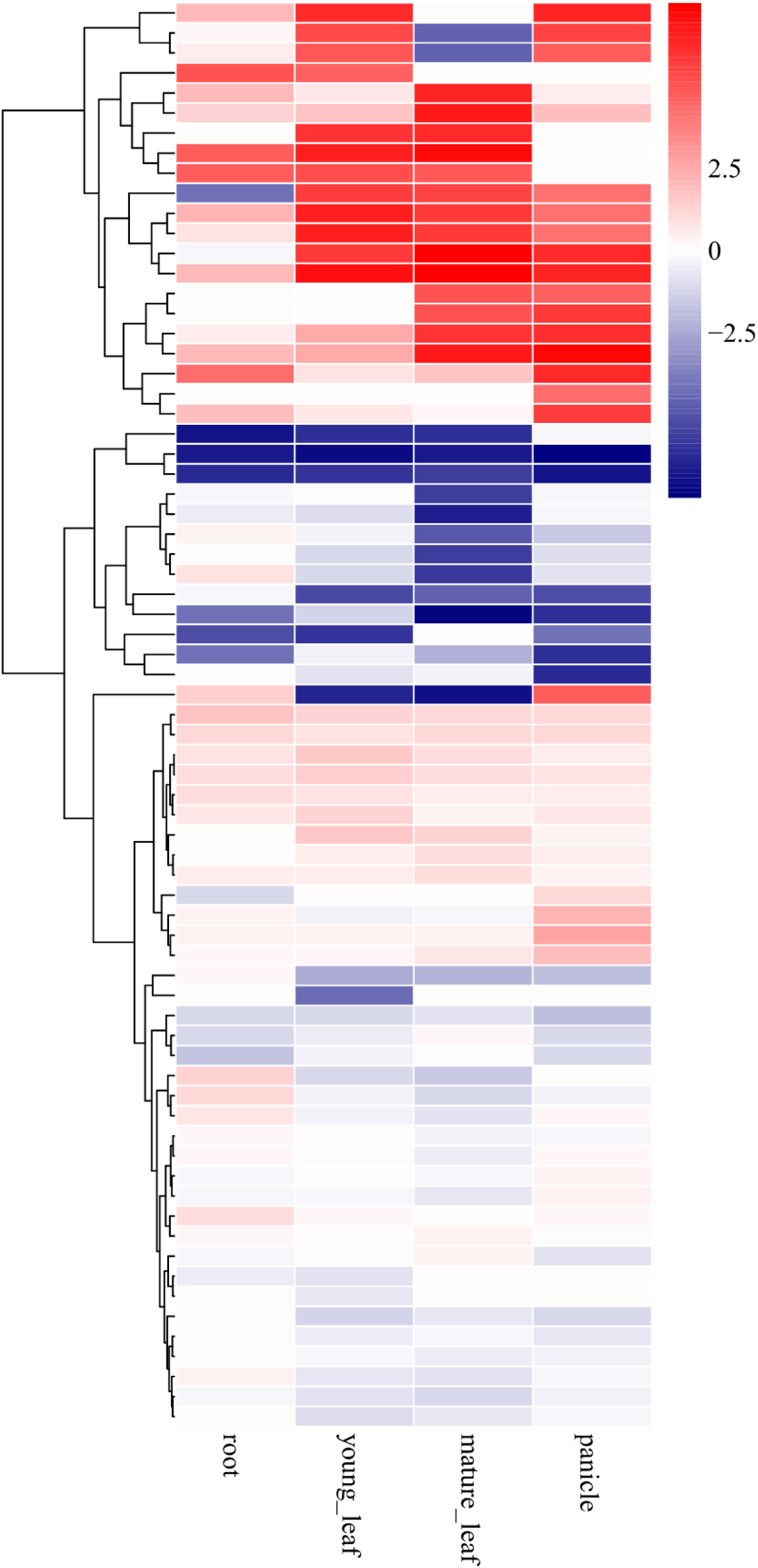
Heatmap of expression ratios for Minghui 63-specific *NBS-LRR* gene pairs. For each duplicate pair, the ratios show the tissue-specific expression level of each gene relative to its duplicate.

**Supplemental Figure 16.**
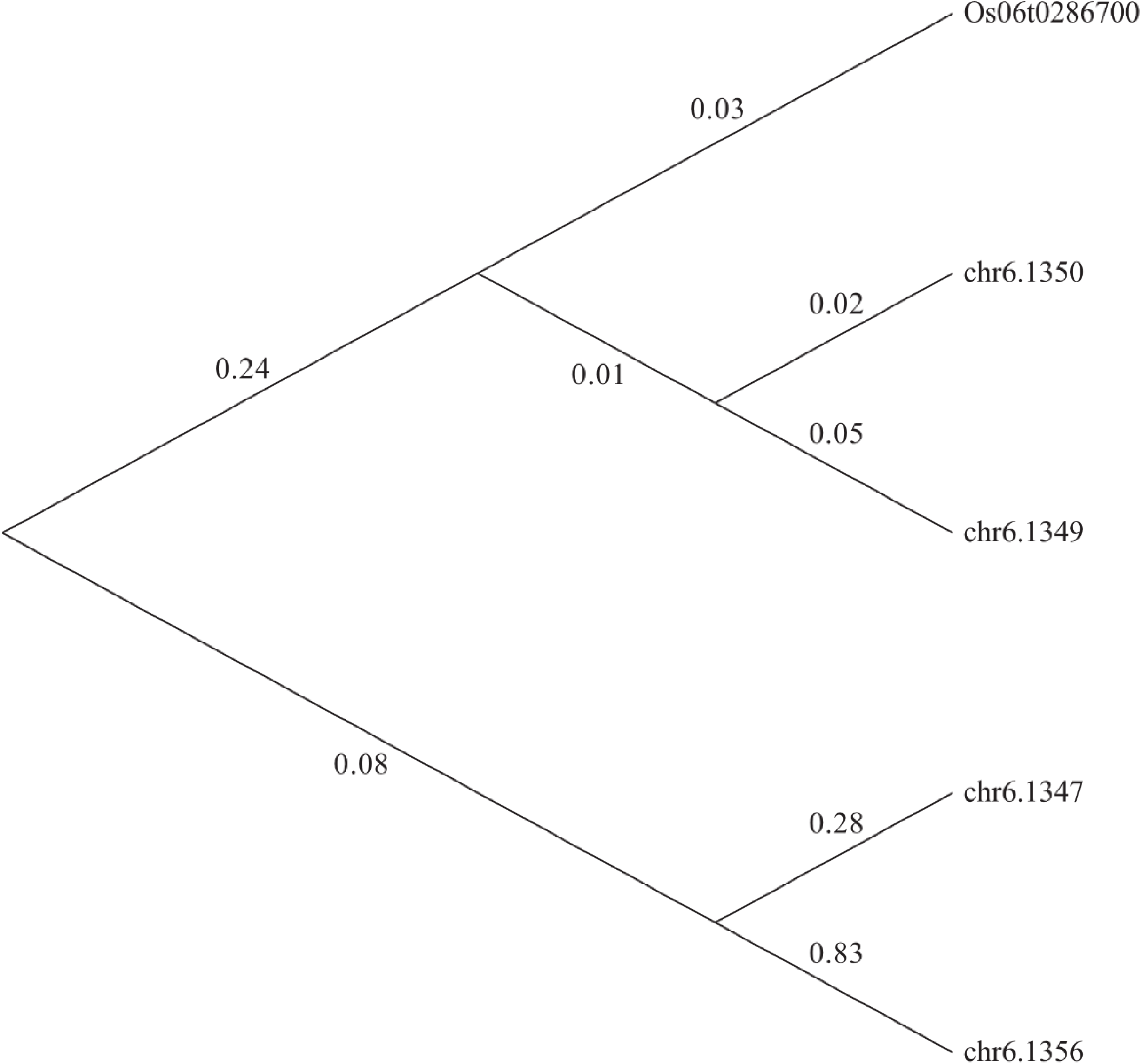
Phylogenetic analysis of rice blast-resistance *Piz* genes. The maximum-likelihood phylogenetic trees were constructed using RAxML (version 8.0.19).

## SUPPLEMENTARY TABLES

**Supplemental Table 1.**
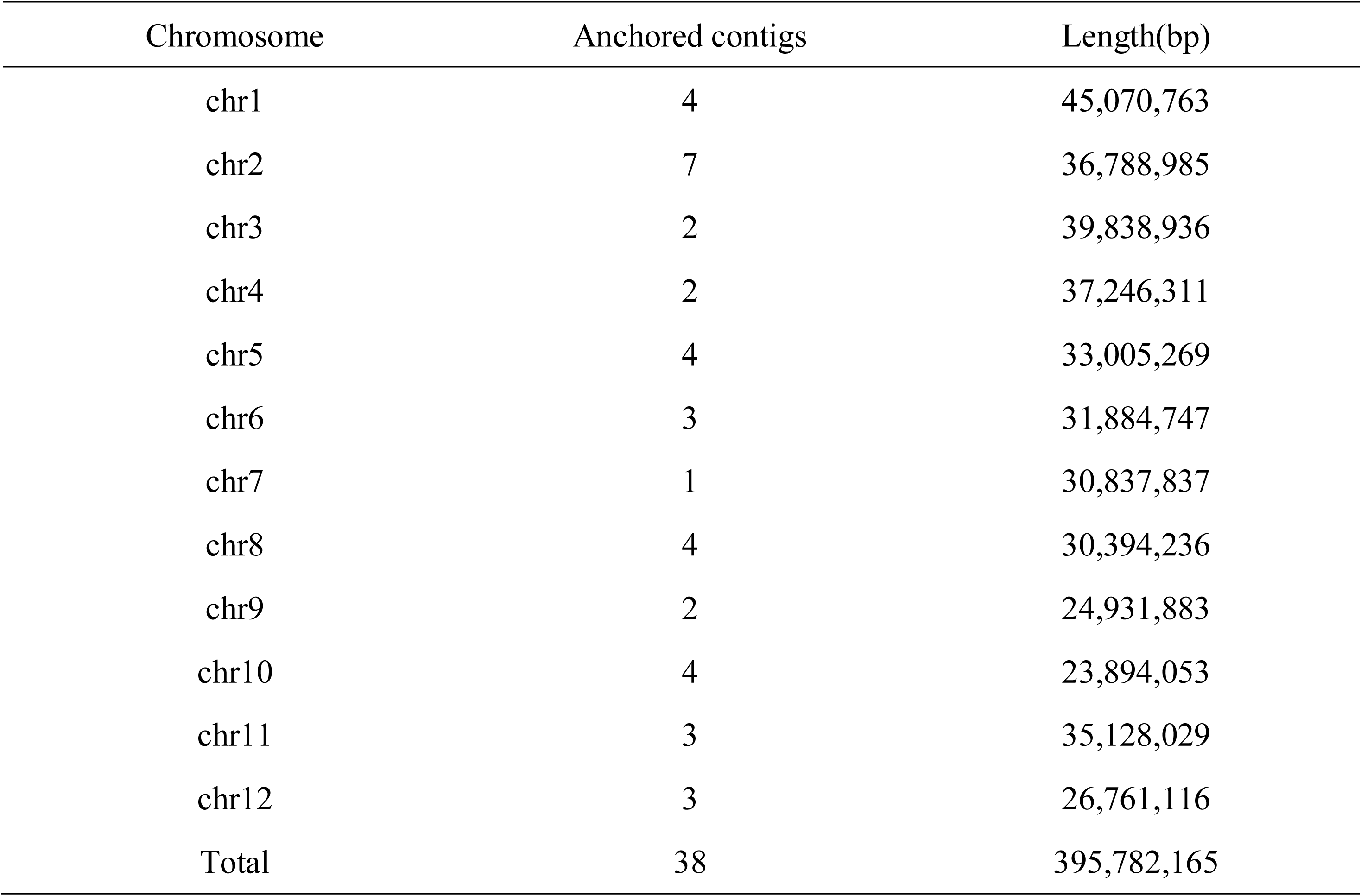
Statistic of Minghui 63 genome assembly.

**Supplemental Table 2.**
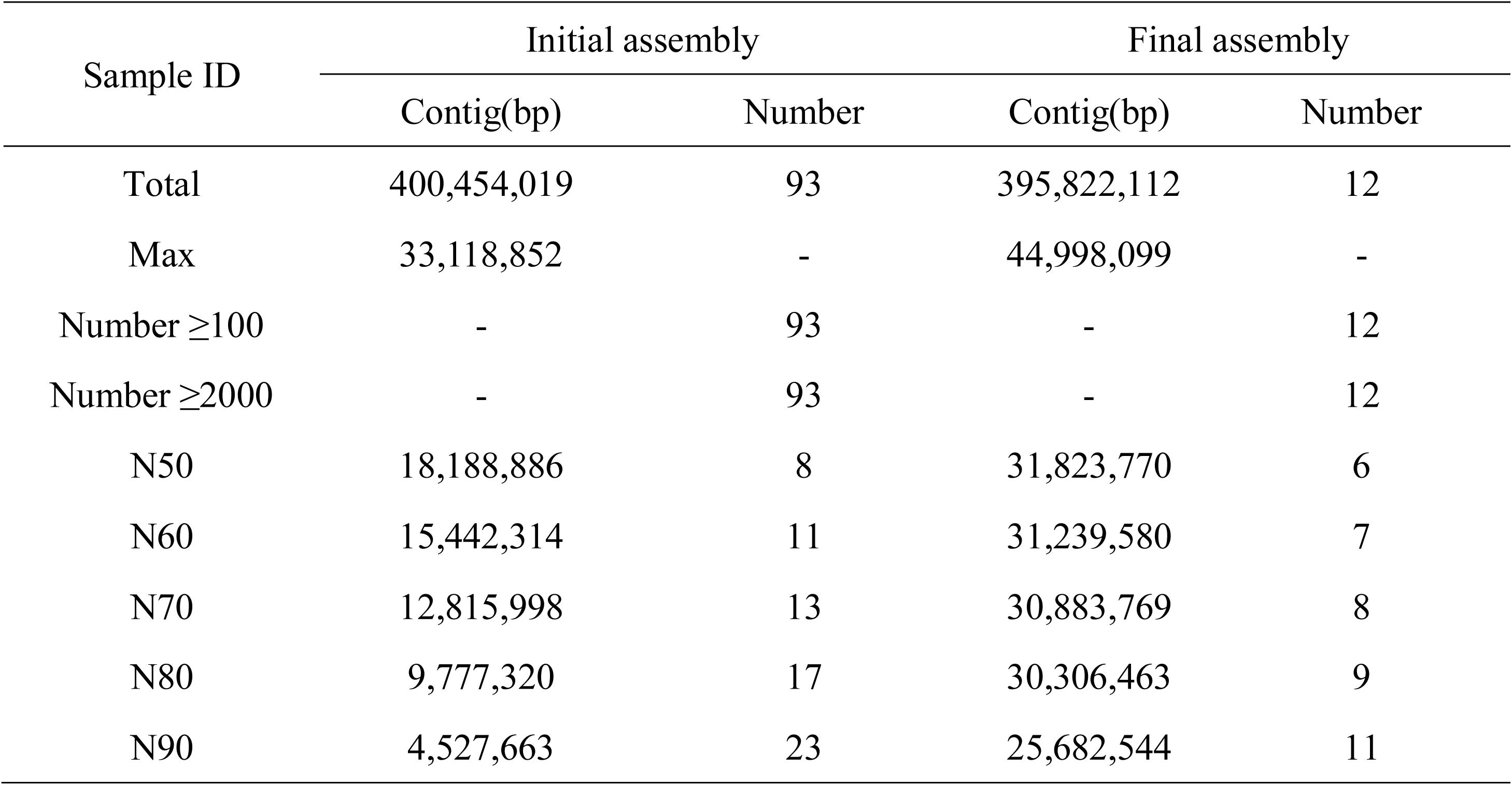
Statistic of Minghui 63 genome chromosome anchoring.

**Supplemental Table 3.** The alignment between 24 gaps and MH63RS2.

| Gap ID | Chr ID | Gap length (bp) | Alignment coverage (%) | Alignment length |
| --- | --- | --- | --- | --- |
| chr1-013-00 | Chr01 | 26,449 | 100 | 26,449 |
| chr1-015-00 | Chr01 | 41,095 | 100 | 41,095 |
| chr2-008-00 | Chr02 | 227,150 | 100 | 227,150 |
| chr2-009-00 | Chr08 | 346,200 | 88.62 | 306,793 |
| chr2-010-00 | Chr02 | 332,102 | 100 | 332,102 |
| chr2-011-00 | Chr02 | 172,627 | 100 | 172,627 |
| chr2-012-00 | Chr02 | 56,472 | 100 | 56,472 |
| chr3-007-00 | Chr03 | 72,573 | 99.99 | 72,569 |
| chr4-006-00 | Chr04 | 66,912 | 100 | 66,912 |
| chr5-003-00 | Chr03 | 87,199 | 100 | 87,199 |
| chr5-004-00 | Chr05 | 84,405 | 100 | 84,405 |
| chr5-005-00 | Chr10 | 616,719 | 87.85 | 541,779 |
| chr6-001-00 | Chr06 | 608,305 | 100 | 608,304 |
| chr6-002-00 | Chr06 | 383,428 | 52.96 | 203,067 |
| chr8-017-00 | Chr08 | 97,121 | 100 | 97,121 |
| chr8-018-00 | Chr08 | 2,177 | 100 | 2,177 |
| chr8-019-00 | Chr08 | 21,090 | 100 | 21,090 |
| chr9-016-00 | Chr09 | 386,058 | 100 | 386,058 |
| chr10-024-00 | Chr10 | 648,325 | 100 | 648,321 |
| chr10-025-00 | Chr10 | 520,537 | 100 | 520,537 |
| chr11-022-00 | Chr11 | 36,960 | 100 | 36,959 |
| chr11-023-00 | Chr11 | 77,603 | 100 | 77,603 |
| chr12-020-00 | Chr12 | 127,827 | 100 | 127,827 |
| chr12-021-00 | Chr12 | 72,384 | 100 | 72,383 |

**Supplementary Table 4.** Statistics of paired-end reads mapping.

|  |  | Percent (%) |
| --- | --- | --- |
| Reads | Mapping rate (%) | 99.34 |
| Genome | Average sequencing depth | 67.81 |
|  | Coverage (%) | 98.98 |
|  | Coverage at least 4X (%) | 97.10 |
|  | Coverage at least 10X (%) | 93.78 |

**Supplementary Table 5.** Assessment of Minghui 63 genome using 2045 Oryza rufipogon W1943 full-length cDNA sequences (http://server.ncgr.ac.cn/ricd/dym/ftp.php).

| Dataset | Number | with >90% sequence in<br>one scaffold |  | with >50% sequence<br>in one scaffold |  |
| --- | --- | --- | --- | --- | --- |
|  |  | Number | Percent | Number | Percent |
| >0bp | 2,045 | 1,880 | 91.93 | 1,971 | 96.38 |
| >200bp | 2,043 | 1,878 | 91.92 | 1,969 | 96.38 |
| >500bp | 1,872 | 1,724 | 92.09 | 1,806 | 96.47 |
| >1000bp | 620 | 583 | 94.03 | 606 | 97.74 |

**Supplementary Table 6.**
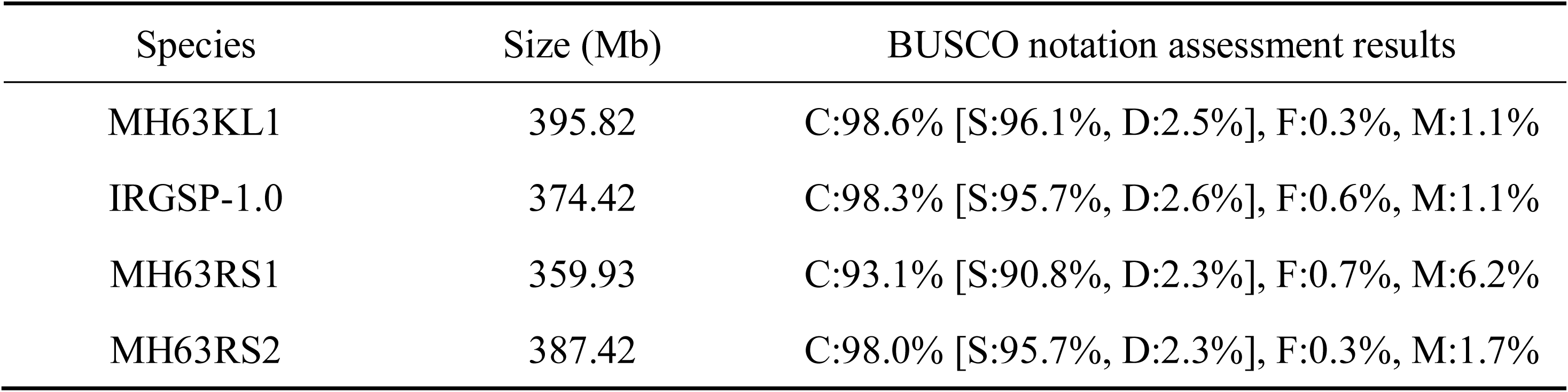
Assessment the gene coverage rate using BUSCO.

| Species | Size (Mb) | BUSCO notation assessment results |
| --- | --- | --- |
| MH63KL1 | 395.82 | C:98.6% [S:96.1%, D:2.5%], F:0.3%, M:1.1% |
| IRGSP-1.0 | 374.42 | C:98.3% [S:95.7%, D:2.6%], F:0.6%, M:1.1% |
| MH63RS1 | 359.93 | C:93.1% [S:90.8%, D:2.3%], F:0.7%, M:6.2% |
| MH63RS2 | 387.42 | C:98.0% [S:95.7%, D:2.3%], F:0.3%, M:1.7% |

**Supplemental Table 7.**
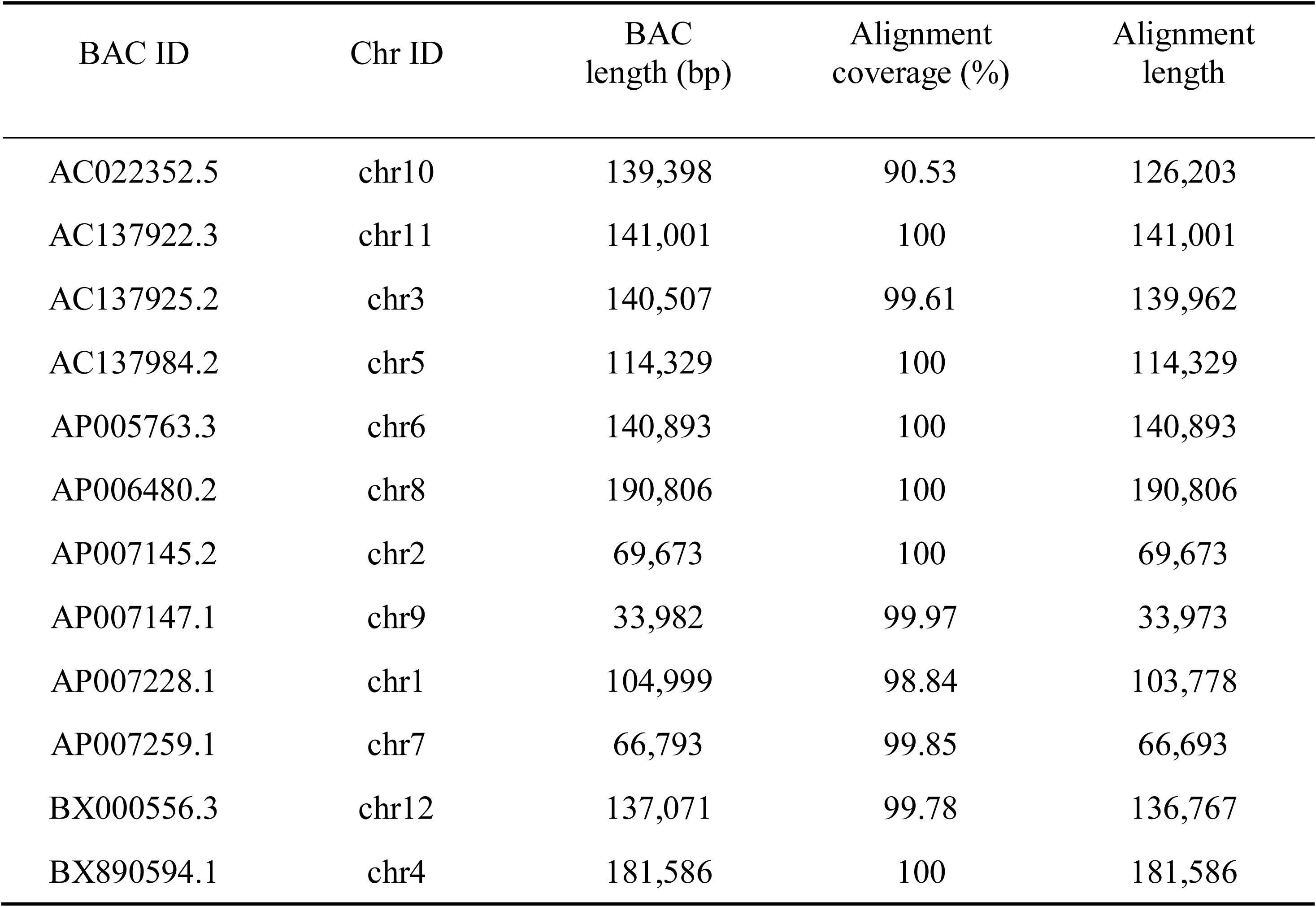
Assessment of the Minghui 63 genome assembly using the whole sequence of 12 BACs of centromeric regions.

**Supplemental Table 8.** The prediction of repeat elements in Minghui 63 genome.

| Type | Repeat Size (bp) | Percent (%) |
| --- | --- | --- |
| TRF | 17,612,333 | 4.45 |
| RepeatMasker | 148,491,480 | 37.51 |
| RepeatProteinMask | 48,979,668 | 12.37 |
| Total | 163,341,094 | 41.27 |

The RepeatMasker and RepeatProteinMask were used to identify TEs. The TRF was used to discover tandem repeats. The Total was integrated of all repeat elements without redundancy.

**Supplemental Table 9.** Categories of TEs predicted in Minghui 63 genome.

| Type | Repeatmasker |  | TE Proteins |  | Combined TEs |  |
| --- | --- | --- | --- | --- | --- | --- |
|  | Length (bp) | % in Genome | Length (bp) | % in Genome | Length (bp) | % in Genome |
| DNA | 21,016,884 | 5.31 | 9,536,305 | 2.41 | 26,015,497 | 6.57 |
| LINE | 2,154,546 | 0.54 | 4,320,725 | 1.09 | 5,832,459 | 1.47 |
| SINE | 112,631 | 0.03 | 0 | 0 | 112,631 | 0.03 |
| LTR | 109,277,567 | 27.61 | 35,229,749 | 8.90 | 112,129,584 | 28.33 |
| Unknown | 2,166,311 | 0.55 | 25,047 | 0.01 | 2,191,358 | 0.55 |
| Total | 148,491,480 | 37.51 | 48,979,668 | 12.37 | 157,157,235 | 39.70 |

**Supplemental Table 10.**
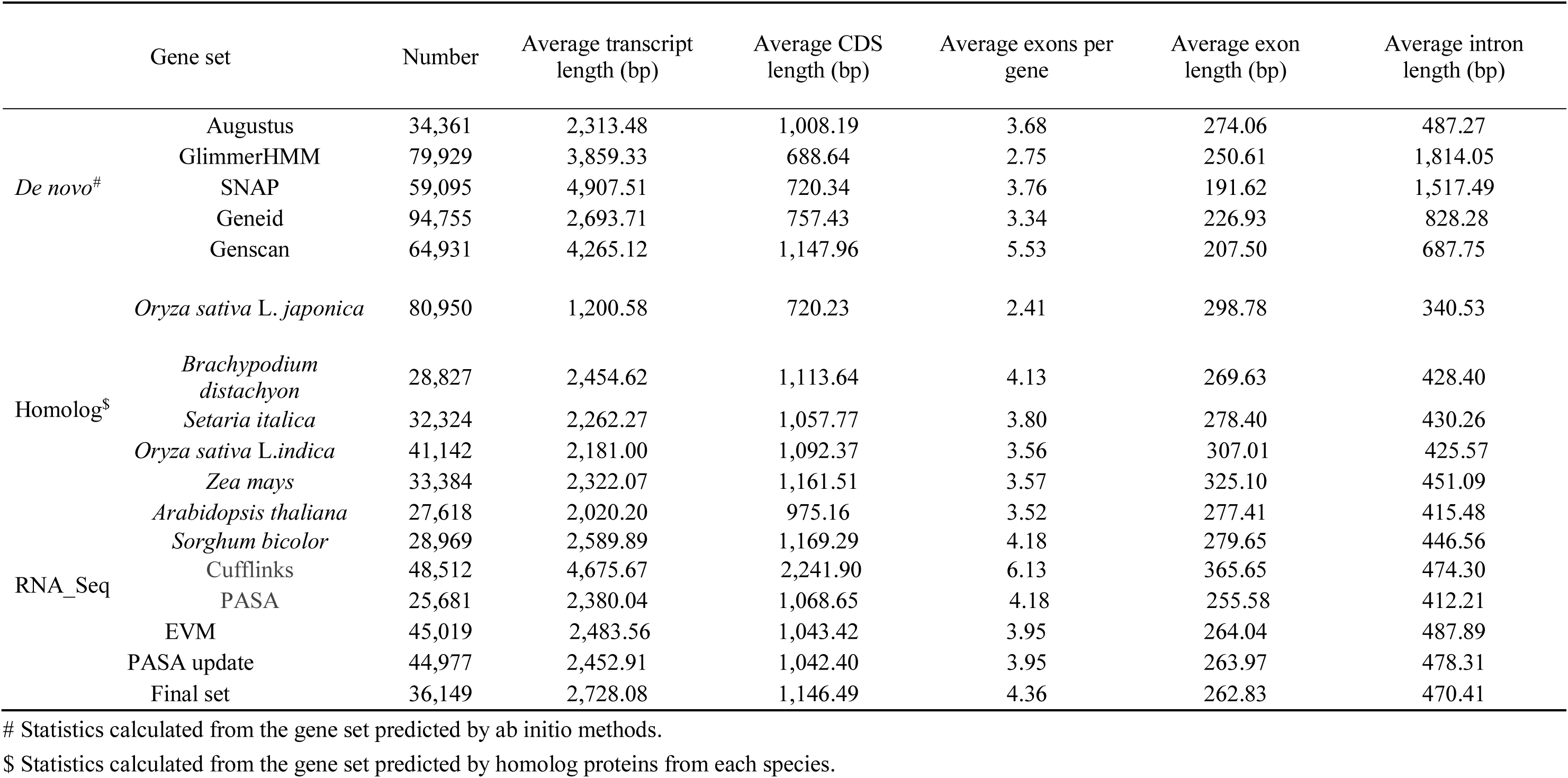
General statistics of predicted protein-coding genes.

**Supplemental Table 11.** General statistics of protein-coding genes of Minghui 63 and homolog species.

| Species | Number | Average transcript length (bp) | Average CDS length (bp) | Average exons per gene | Average exon length (bp) | Average intron length (bp) |
| --- | --- | --- | --- | --- | --- | --- |
| *Oryza sativa* L.indica | 36,149 | 2,728.08 | 1,146.49 | 4.36 | 262.83 | 470.41 |
| *Oryza sativa* L. japonica | 35,311 | 2,200.75 | 1,008.79 | 3.83 | 263.56 | 421.56 |
| *Brachypodium distachyon* | 34,014 | 2,524.76 | 1,140.59 | 4.46 | 255.70 | 399.98 |
| *Setaria italica* | 33,664 | 2,308.30 | 1,165.43 | 4.54 | 256.45 | 322.44 |
| *Zea mays* | 39,235 | 3,485.39 | 1,183.48 | 4.93 | 239.91 | 585.29 |
| *Sorghum bicolor* | 33,878 | 2,771.31 | 1,169.67 | 4.53 | 257.97 | 453.19 |

**Supplemental Table 12.** General statistics of functional annotation of protein-coding genes.

|  | Number | Percent (%) |
| --- | --- | --- |
| Total | 36,149 | - |
| Swissprot | 22,729 | 62.9 |
| NR | 34,965 | 96.7 |
| KEGG | 21,520 | 59.5 |
| InterPro | 35,650 | 98.6 |
| Pfam | 24,194 | 66.9 |
| GO | 32,381 | 89.6 |
| Annotated | 36,062 | 99.8 |
| Unannotated | 87 | 0.2 |

**Supplemental Table 13.** CentO (155 bp satellite DNA) units in Minghui 63 genome.

| Chromosome | CentO sequence location |  | Total no. of units | Total length (bp) |
| --- | --- | --- | --- | --- |
|  | Start | End |  |  |
| chr1 | 17,510,384 | 18,355,493 | 1,111 | 845,110 |
| chr2 | 13,502,971 | 13,829,567 | 683 | 326,597 |
| chr3 | 21,134,162 | 21,646,956 | 124 | 512,795 |
| chr4 | 9,549,585 | 9,697,276 | 307 | 147,692 |
| chr5 | 12,537,089 | 12,656,508 | 707 | 119,420 |
| chr6 | 15,460,307 | 15,580,164 | 322 | 119,858 |
| chr7 | 12,448,237 | 12,450,250 | 13 | 2,014 |
| chr7 | 13,139,270 | 13,143,534 | 27 | 4,265 |
| chr8 | 13,481,133 | 13,583,132 | 183 | 102,000 |
| chr8 | 14,466,603 | 14,536,492 | 13 | 69,890 |
| chr9 | 2,612,761 | 2,654,031 | 56 | 41,271 |
| chr9 | 3,105,486 | 3,289,248 | 149 | 183,763 |
| chr10 | 9,456,089 | 9,495,536 | 5 | 39,448 |
| chr11 | 13,789,153 | 13,806,601 | 59 | 17,449 |
| chr12 | 10,570,208 | 11,003,912 | 549 | 433,705 |
| Total | - | - | 4,308 | 2,965,277 |

**Supplemental Table 14.** Statistic distribution of SDs on chromosomes. The overlapped blocks are merged using bedtools.

| Chromosome | SDs number | SD Length (bp) | Chr length (bp) | percentage (%) |
| --- | --- | --- | --- | --- |
| chr1 | 2,452 | 7,819,396 | 44,998,099 | 17.38 |
| chr2 | 1,992 | 6,831,623 | 36,790,212 | 18.57 |
| chr3 | 2,025 | 6,775,221 | 39,918,125 | 16.97 |
| chr4 | 1,934 | 9,505,882 | 37,315,983 | 25.47 |
| chr5 | 1,762 | 6,279,285 | 31,239,580 | 20.10 |
| chr6 | 1,808 | 6,999,350 | 31,823,770 | 21.99 |
| chr7 | 1,933 | 6,942,480 | 30,883,769 | 22.48 |
| chr8 | 1,778 | 7,157,808 | 30,306,463 | 23.62 |
| chr9 | 1,467 | 5,540,613 | 24,951,677 | 22.21 |
| chr10 | 1,622 | 6,720,233 | 25,682,544 | 26.17 |
| chr11 | 2,103 | 11,236,816 | 35,116,109 | 32.00 |
| chr12 | 1,571 | 7,875,604 | 26,795,781 | 29.39 |
| Total | 22,447 | 89,684,311 | 395,822,112 | 22.66 |

**Supplemental Table 15.** The KEGG enrichment analysis of the subfunctional gene pairs.

| MapID | MapTitle | Pvalue | Number |
| --- | --- | --- | --- |
| map00908 | Zeatin biosynthesis | 9.01E-07 | 7 |
| map00460 | Cyanoamino acid metabolism | 7.40E-05 | 9 |
| map00960 | Tropane, piperidine and pyridine alkaloid biosynthesis | 3.27E-04 | 6 |
| map00260 | Glycine, serine and threonine metabolism | 4.82E-04 | 8 |
| map00950 | Isoquinoline alkaloid biosynthesis | 6.22E-04 | 6 |
| map00073 | Cutin, suberine and wax biosynthesis | 1.30E-03 | 6 |
| map00480 | Glutathione metabolism | 1.31E-03 | 8 |
| map00942 | Anthocyanin biosynthesis | 1.48E-03 | 4 |
| map00410 | beta-Alanine metabolism | 1.53E-03 | 6 |
| map00906 | Carotenoid biosynthesis | 1.89E-03 | 6 |
| map00360 | Phenylalanine metabolism | 2.09E-03 | 6 |
| map00350 | Tyrosine metabolism | 2.80E-03 | 6 |
| map00903 | Limonene and pinene degradation | 4.70E-03 | 4 |
| map00905 | Brassinosteroid biosynthesis | 8.27E-03 | 3 |
| map00565 | Ether lipid metabolism | 1.27E-02 | 4 |
| map00062 | Fatty acid elongation | 1.33E-02 | 4 |
| map00630 | Glyoxylate and dicarboxylate metabolism | 4.48E-02 | 4 |
| map00790 | Folate biosynthesis | 4.82E-02 | 2 |

**Supplemental Table 16.** General statistics of intact LTR-RTs.

| Species | Total Length<br>(bp) | Total<br>Number | Max<br>length (bp) | Min<br>length (bp) | Average<br>length (bp) |
| --- | --- | --- | --- | --- | --- |
| MH63KL1 | 53,645,453 | 5,516 | 25,673 | 1,165 | 9,725.43 |
| IRGSP-1.0 | 43,495,559 | 4,706 | 24,598 | 1,217 | 9,242.58 |

**Supplemental Table 17.** The classification of LTR-RTs.

| MH63KL1 |  | IRGSP-1.0 |  |
| --- | --- | --- | --- |
| family | number | family | number |
| Del | 1,911 | Tat | 1,223 |
| Tat | 1,057 | Del | 923 |
| Tork | 373 | Tork | 455 |
| Crm | 319 | Crm | 238 |
| Sire | 187 | Sire | 231 |
| Reina | 139 | Reina | 143 |
| Oryco | 131 | Retrofit | 129 |
| Retrofit | 117 | Oryco | 127 |
| Athila | 91 | Athila | 114 |
| Copia | 31 | Copia | 34 |

**Supplemental Table 18.** General statistics of *NBS-LRR* genes.

| **NBS type** | **MH63KL1** | **IRGSP-1.0** |
| --- | --- | --- |
| TIR-NBS | 2 | 1 |
| CC_NBS_LRR | 25 | 8 |
| CC_NBS | 92 | 67 |
| NBS_LRR | 77 | 51 |
| NBS | 291 | 222 |
| Total | 487 | 349 |

**Supplemental Table 19.** The distribution of *NBS-LRR* genes on each chromosome.

| chr | MH63KL1 | IRGSP-1.0 |
| --- | --- | --- |
| chr1 | 44 | 35 |
| chr2 | 30 | 20 |
| chr3 | 19 | 11 |
| chr4 | 27 | 15 |
| chr5 | 23 | 12 |
| chr6 | 31 | 25 |
| chr7 | 39 | 23 |
| chr8 | 47 | 35 |
| chr9 | 26 | 16 |
| chr10 | 33 | 24 |
| chr11 | 123 | 98 |
| chr12 | 45 | 35 |

**Supplemental Table 20.**
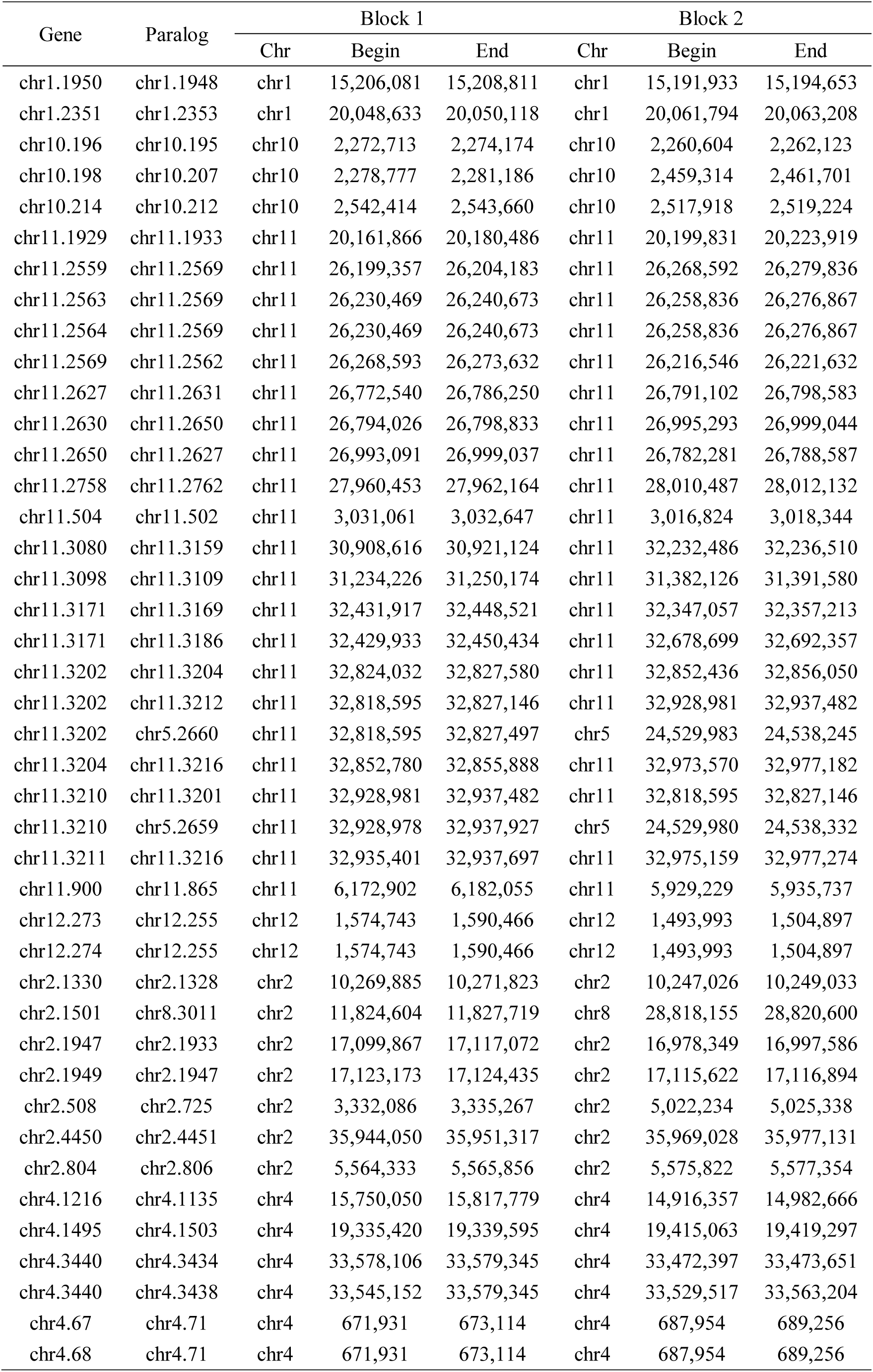
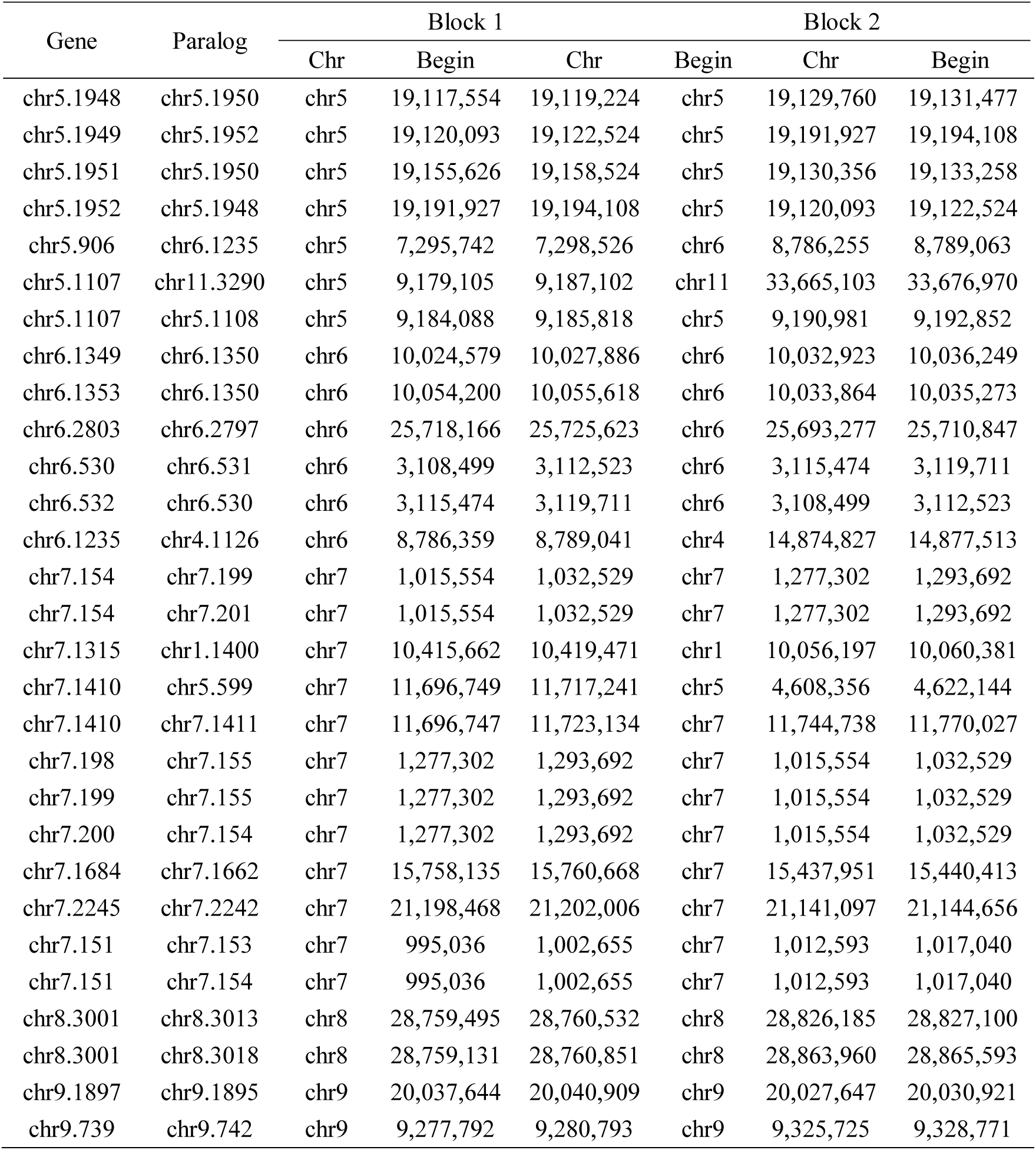
SDs collinear block pairs of Minghui 63-specific *NBS-LRR* genes.

**Supplemental Table 21.**
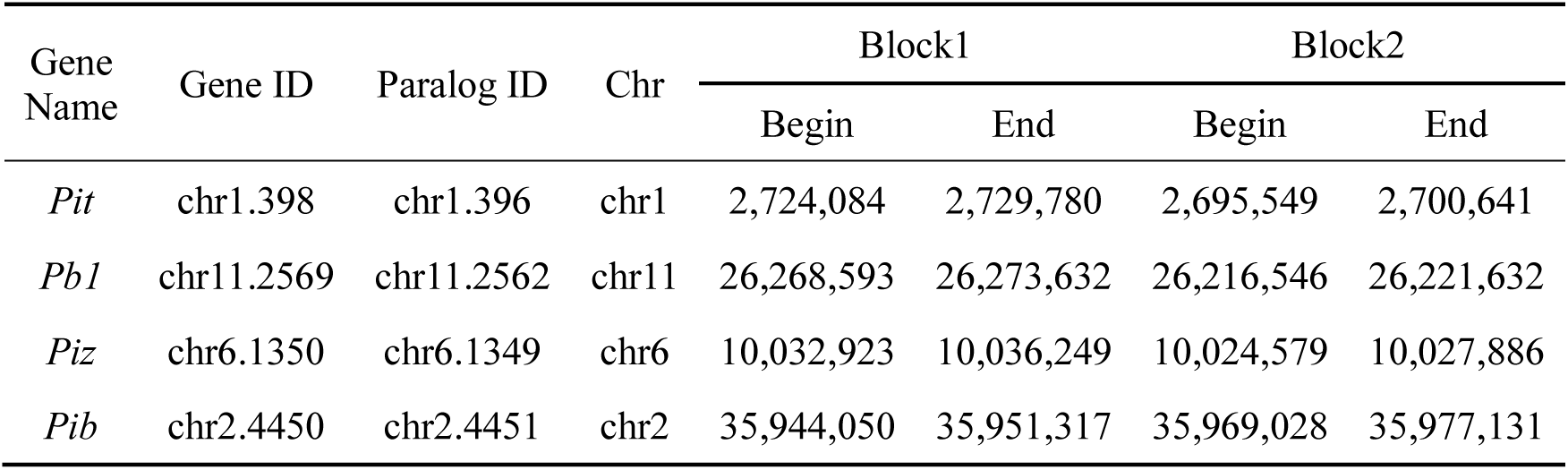
SDs collinear block pairs of four rice blast resistance genes.

**Supplemental Table 22.** SDs collinear block pairs of *cis*-zeatin-*O*-glucosyltransferase genes.

| Gene ID | Paralog ID | Chr | Block1 |  | Block2 |  |
| --- | --- | --- | --- | --- | --- | --- |
|  |  |  | Begin | End | Begin | End |
| chr4.2981 | chr4.2983 | chr4 | 30,071,801 | 30,073,214 | 30,093,424 | 30,094,837 |
| chr4.2909 | chr4.2910 | chr4 | 29,601,285 | 29,603,206 | 29,606,146 | 29,607,947 |

